# Targeting RUNX1 in Macrophages Facilitates Cardiac Recovery

**DOI:** 10.1101/2025.07.25.665779

**Authors:** Junedh M. Amrute, Ada Zhu, Yun-Ling Pai, Melissa Hector-Greene, Yuqian An, Kenji Rowell Kim, Maya U. Sheth, Arun Padmanabhan, Clara Youngna Lee, Tracy Yamawaki, Florian Sicklinger, Niklas Hartman, Andrea Bredemeyer, Chang Jie Mick Lee, Vee Xu, Lauren Bell, Tyler Harmon, Haewon Shin, Alekhya Parvathaneni, Lei Liu, Amal K. Dutta, Danielle Pruitt, Jose Barreda, Jing Chen, Urvi Nikhil Shroff, Rangarajan Nadadur, Jess Nigro, Carla Weinheimer, Atilla Kovacs, Jixin Cui, Chen Wang, Chi-Ming Li, Daniel Kreisel, Yongjian Liu, Roger S-Y Foo, Rebekka K Schneider, Jesse M. Engreitz, Douglas L. Mann, Ingrid Rulifson, Simon Jackson, Brandon Ason, Rafael Kramann, Stavros G. Drakos, Florian Leuschner, Michael Alexanian, Kory J. Lavine

**Affiliations:** Center for Cardiovascular Research, Division of Cardiology, Department of Medicine, Washington University School of Medicine, Saint Louis, MO, 63110, USA; Amgen, 750 Gateway Blvd, South San Francisco, CA, 94080, USA; Gladstone Institutes, San Francisco, 94080, CA, USA; Roddenberry Center for Stem Cell Biology at Gladstone Institutes, San Francisco, 94080, CA, USA; Department of Genetics, Stanford University School of Medicine, Stanford, CA, USA; Basic Sciences and Engineering Initiative, Betty Irene Moore Children’s Heart Center, Lucile Packard Children’s Hospital, Stanford, CA, USA; Department of Bioengineering, Stanford University, Stanford, CA, USA; Department of Medicine, Division of Cardiology, University of California, San Francisco, San Francisco, CA, USA; Chan Zuckerberg Biohub San Francisco, San Francisco, CA, USA; Department of Cardiology, University Hospital Heidelberg, Heidelberg, Germany; German Centre for Cardiovascular Research (DZHK), Partner Site Heidelberg, Heidelberg, Germany; Cardiovascular Metabolic Disease Translational Research Programme, National University Health System, Centre for Translational Medicine, 14 Medical Drive, Singapore 117599, Singapore; Institute of Molecular and Cell Biology, 61 Biopolis Drive, Singapore 138673, Singapore; Department of Pathology and Immunology, Washington University School of Medicine, Saint Louis, MO, 63110, USA; Division of Cardiothoracic Surgery, Department of Surgery, Washington University School of Medicine, Saint Louis, 63110, MO, USA; Mallinckrodt Institute of Radiology, Washington University School of Medicine, Saint Louis, MO, 63110, USA; Department of Developmental Biology, Erasmus Medical Center Cancer Institute, Rotterdam, Netherlands; Oncode Institute, Erasmus MC, Rotterdam, Netherlands; Department of Cell and Tumor Biology, Faculty of Medicine, RWTH Aachen University, Aachen, Germany; The Novo Nordisk Foundation Center for Genomic Mechanisms of Disease, Broad Institute, Cambridge, MA, USA; Stanford Cardiovascular Institute, Stanford University, Stanford, CA 94305; Gene Regulation Observatory, Broad Institute of MIT and Harvard, Cambridge, MA 02139, USA; Institute of Experimental Medicine and Systems Biology and Division of Nephrology, RWTH Aachen University, Aachen, Germany; Department of Internal Medicine, Erasmus Medical Center, Rotterdam, The Netherlands; Division of Cardiovascular Medicine & Nora Eccles Harrison Cardiovascular Research and Training Institute, University of Utah Health & School of Medicine, Salt Lake City, UT; Department of Pediatrics, University of California, San Francisco, San Francisco, CA, USA; Department of Developmental Biology, Washington University School of Medicine, Saint Louis, MO, 63110, USA

**Keywords:** Organ recovery, heart failure, inflammation, Runt-related transcription factor 1 (RUNX1), single cell multiome, chromatin, CRISPR

## Abstract

Despite advances in disease treatment, our understanding of how damaged organs recover and the mechanisms governing this process remain poorly defined. Here, we mapped the transcriptional and regulatory landscape of human cardiac recovery using single cell multiomics. Macrophages emerged as the most reprogrammed cell type. Deep learning identified the transcription factor RUNX1 as a key regulator of this process. Macrophage-specific *Runx1* deletion recapitulated the human cardiac recovery phenotype in a chronic heart failure model. *Runx1* deletion reprogrammed macrophages to a reparative phenotype, reduced fibrosis, and promoted cardiomyocyte adaptation. RUNX1 chromatin profiling revealed a conserved regulon that diminished during recovery. Mechanistically, the epigenetic reader BRD4 controlled *Runx1* expression in macrophages. Chromatin activity mapping, combined with CRISPR perturbations, identified the precise regulatory element governing *Runx1* expression. Therapeutically, small molecule Runx1 inhibition was sufficient to promote cardiac recovery. Our findings uncover a druggable RUNX1 epigenetic mechanism that orchestrates recovery of heart function.

## Introduction

Across a range of chronic diseases, damaged organs can regain function^1–4^. However, the molecular mechanisms that govern this process remain poorly defined. Understanding biological principles of how organs recover from pathological states holds promise to unlock new therapeutic strategies. Heart failure (HF) represents an ideal disease entity to study this phenomenon^1,3–11^. Affecting over 60 million people worldwide, HF remains a leading cause of death globally despite modern treatment approaches^12–15^. In concert with medical therapy for heart failure, mechanical unloading of the failing heart using left ventricular assist devices (LVADs) remarkably leads to myocardial recovery in a small proportion of patients^1–11,16–21^. However, the molecular and cellular mechanisms that distinguish patients who recover from those who will experience disease progression remain unknown. Defining the gene programs and cell states that underlie cardiac recovery may open new avenues for restoring heart function in individuals with advanced and refractory disease.

Recent studies have used single-cell and single-nucleus technologies to delineate the cellular heterogeneity of failing human hearts^6,22–30^, uncovering maladaptive cell states in cardiomyocytes, fibroblasts, endothelial cells, and immune cells. While significant progress has been made in understanding disease progression, the molecular basis of cardiac recovery remains largely unknown. In the heart, macrophages have emerged as key regulators of inflammation and tissue remodeling^25,31–36^, yet their role in orchestrating cardiac recovery remains unknown.

Here, we leveraged single-nucleus gene expression and chromatin profiling from non-failing donors, patients with end-stage HF, and those who recovered post-LVAD implantation^6,25^ to uncover gene and chromatin programs associated with myocardial recovery. Among major cell types, macrophages exhibit the strongest recovery-associated transcriptional signature, characterized by suppression of pro-inflammatory gene programs and emergence of glucocorticoid-responsive states. Using deep learning and gene regulatory network modeling, we identified the transcription factor Runt-related transcription factor 1 (*RUNX1*) as a central regulator of a pathologic macrophage program negatively associated with recovery. In silico perturbation^37^ of *RUNX1* reprogrammed macrophages toward a recovery-like state, supporting its potential as a gene regulatory network-correcting target.

To validate these findings, we employed a murine model of chronic heart failure^38^. Loss of *Runx1* in cardiac macrophages resulted in functional, physiological, and molecular recovery, with restoration of a transcriptional signature seen in human recovery. Mechanistically, *Runx1* deletion suppressed inflammatory macrophage states and reprogrammed fibroblasts away from pro-fibrotic phenotypes. These effects were accompanied by the induction of conserved recovery-associated genes, including *Zbtb16* and *Tsc22d3*, across multiple cell types in human and mice. We further show that *Runx1* expression in macrophages is dynamically regulated by the chromatin reader bromodomain-containing protein 4 (*BRD4*) in response to cardiac stress. Using murine BRD4 knockout models^31^, *in vivo* BRD4 CUT&RUN mapping, and CRISPR interference (CRISPRi) experiments we defined the exact molecular mechanisms governing stress-induced *Runx1* expression. We then performed in vivo gene expression and chromatin accessibility profiling in *Runx1* knockout animals, coupled with CUT&RUN, to construct a macrophage-specific RUNX1 regulon that was suppressed in human myocardial recovery. Finally, pharmacologic disruption of the *RUNX1* network using a small molecule inhibitor^39^ suppressed pro-inflammatory signaling and promoted cardiac recovery in vivo. Together, these findings define a conserved transcriptional program of myocardial recovery orchestrated by macrophages and identify RUNX1 as a pivotal node regulating the transition from heart failure to cardiac recovery.

## Results

### Deep learning in human recovery prioritizes macrophage gene programs

To identify cellular and transcriptional determinants of myocardial recovery in humans, we utilized integrated single-nucleus transcriptomic and chromatin accessibility profiling of hearts from non-failing donors, patients with end-stage heart failure (HF), and patients who recovered function after mechanical unloading^6,25^. Restoration of left ventricular ejection fraction (LVEF)^6,9^ was assessed by echocardiography performed when LVAD support was reduced to the lowest feasible setting. We asked whether cell type specific transcriptional changes were associated with recovery. For each major cardiac cell type, we computed the correlation between gene expression and LVEF across patients, identifying genes whose expression was positively or negatively associated with recovery. Among all major cardiac cell types, macrophages exhibited the greatest degree of dynamic transcriptional correlation with cardiac function, with >150 genes significantly associated with ejection fraction (**Fig. 1A-B** and **Suppl. Fig. 1A**). To further define gene programs specific to myocardial recovery, we performed pairwise pseudobulk differential gene expression analysis across conditions. This approach allowed us to isolate a recovery-specific transcriptional signature—distinct from that of non-failing hearts, end-stage HF, or patients with persistent dysfunction following mechanical unloading (Fig. 1C) (**Fig. 1C**). Notably, we found that macrophages contained the strongest transcriptional signature of recovery (>500 significantly regulated genes) followed by fibroblasts, endothelial cells, endocardial cells, and cardiomyocytes (**Fig. 1C**).

**Figure 1.**
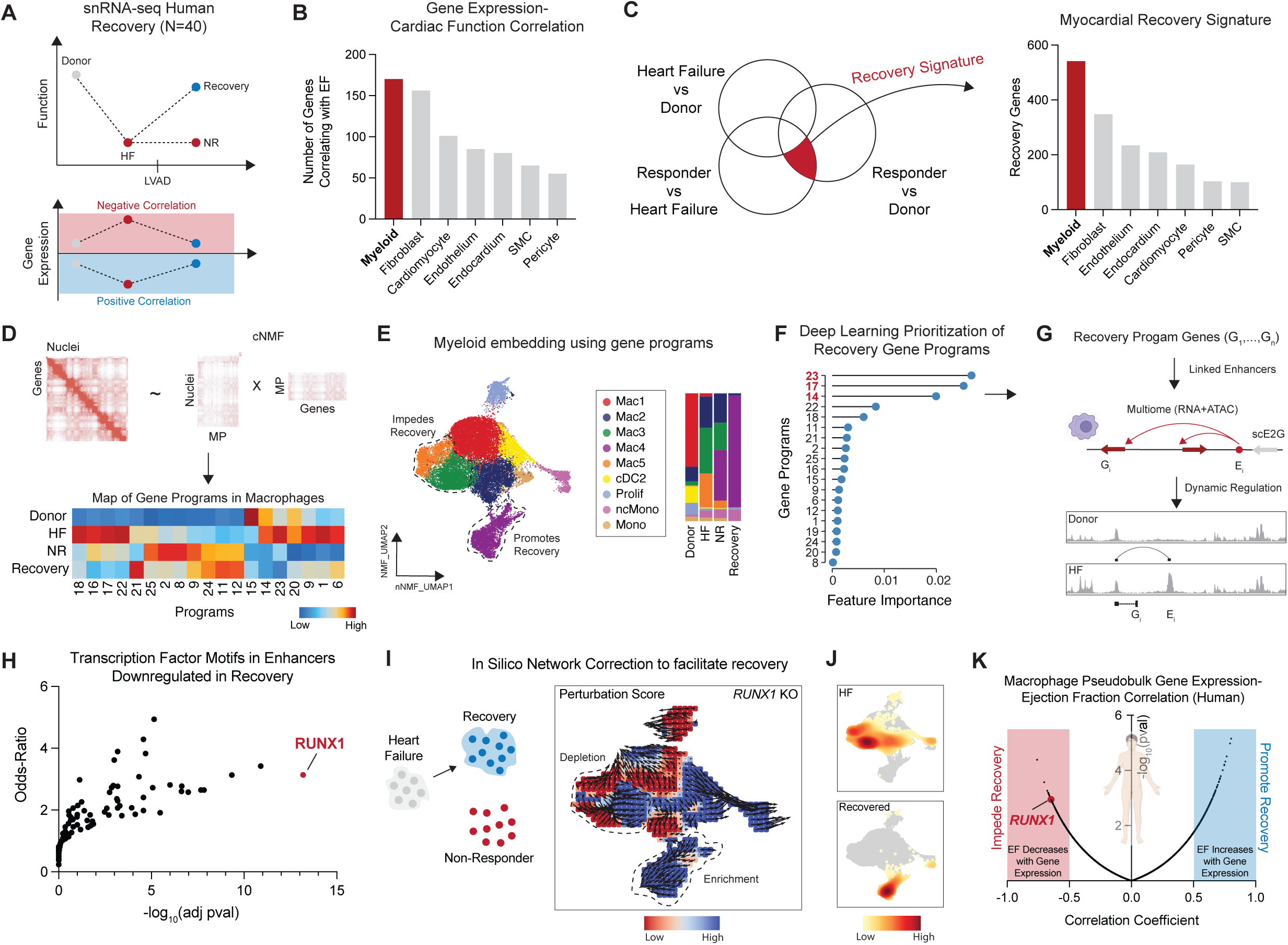
A *RUNX1* gene program in macrophages drives recovery in human hearts. (A) Study design of human myocardial recovery upon mechanical unloading. (B) Number of genes correlating with EF at the pseudobulk level in patients undergoing mechanical unloading and cardiac recovery. (C) Venn diagram depicting intersection of unique recovery associated genes not seen in healthy, failing, or failing mechanically un-loaded hearts (left) and tabulation of unique recovery genes by major cell type (right). (D) cNMF decomposition of macrophages into gene programs (top) and heatmap of program score across patient groups. (E) Myeloid embedding using NMF components for dimensional reduction colored by cell state (left) and relative cell composition by patient group (right). (F) augr predicted feature importance for gene programs predictive of myocardial recovery. (G) Schematic of linking recovery program genes to dynamic enhancers through a scE2G framework in macrophages from human heart failure Multiome (RNA + ATAC) data. (H) Transcription factor motif enrichment in promoters and enhancers linked to recovery program genes highlights significant depletion of RUNX1 motifs in recovery. The adjusted p-value is computed using the Benjamini-Hochberg method for correction for multiple hypothesis testing. The odds-ratio is derived from running the Fisher exact test for many random gene sets to compute a mean rank and standard deviation from the expected rank for each term in the gene set library and calculating a z-score to assess deviation from expected rank. (I) In silico *RUNX1* knock out simulation using CellOracle to shift failing macrophages towards a recovery state. The arrows are vectors pointing towards cell state transitions under the KO condition and colored by perturbation score (computed as the inner dot product of developmental vectors and simulated shifts). (J) Cell density plots of macrophages from HF or recovered individuals. (K) Violin plot of Pearson correlation coefficient between gene expression and ejection fraction at pseudobulk level showing *RUNX1* negatively associates with function. HF = heart failure, NR = non-responder (individuals who do not recover function after mechanical unloading). EF = ejection fraction. cNMF = consensus non-negative matrix factorization. MP = molecular program. scE2G = single cell enhancer-gene.

To resolve the underlying structure of macrophage heterogeneity driving recovery, we applied consensus non-negative matrix factorization (cNMF)^40,41^ and identified twenty-five distinct gene expression programs (**Fig. 1D**). We found that gene programs 21, 11, 12, and 24 were enriched in recovery while gene programs 14, 17, and 23 were most depleted in recovery (**Fig. 1D**). We next projected these transcriptional programs onto a UMAP embedding using nearest-neighbor clustering to define conserved macrophage states across conditions (Fig. 1E) (**Fig. 1E**). This analysis revealed a pro-inflammatory macrophage state (Mac5), marked by high expression of *CCL2, CCL3*, and *RUNX1,* which was enriched in HF and absent in recovered or donor hearts. In contrast, another macrophage state (Mac4), expressing *FKBP5*, *ZBTB16*, and *TSC22D3*, expanded after mechanical unloading and was most prominent in recovered individuals (**Fig. 1E**). Differential gene expression analysis revealed key marker genes associated with each macrophage state (Mac1–Mac5), corresponding to resident-like, antigen-presenting, oxidative stress, glucocorticoid-responsive, and inflammatory states, respectively (**Fig. 1E** and **Suppl. Fig. 1B-D**). To systematically prioritize programs most predictive of recovery, we implemented a deep learning-based model^42^ trained on recovery status, and found that gene programs 23, 17, and 14 were most predictive of recovery (**Fig. 1F**). Next, we performed pathway analysis using genes with program 23 and found enrichment of pro-inflammatory pathways such as NF-kB, IL-6, and JAK/STAT (**Suppl. Fig. 1E**). Together, these findings suggest that pro-inflammatory macrophage states may hinder recovery, while glucocorticoid-responsive macrophages may promote recovery following mechanical unloading.

### In silico perturbation prioritizes RUNX1 as a network-correcting target

To dissect regulatory networks driving recovery gene programs, we adapted a single-cell enhancer-to-gene linkage framework (scE2G)^43,44^ that integrates chromatin accessibility and gene expression from human heart failure multiome (RNA + ATAC) data^25^. We constructed a macrophage specific E2G map and prioritized regulatory elements enriched in failing hearts linked to genes in program 23, the pro-inflammatory macrophage signature that is the strongest negative predicter of recovery (**Fig. 1G**). We then used motif enrichment analysis within the program 23 E2G map to prioritize transcription factors regulating enriched genes within this program (**Fig. 1H**). Notably, we found that *RUNX1* emerged as the top transcriptional regulator, which was also inversely associated with human cardiac recovery (**Fig. 1H**). We then applied *in silico* network correction using CellOracle^37^ to simulate transcription factor perturbation and predict effects on macrophage state composition and acquisition of the recovery phenotype. We simulated in silico *RUNX1* KO and quantified the perturbation score across cell states by calculating the inner product of the developmental pseudotime^45^ and inferring transition vectors representative of how cells shift in the UMAP space. We found that *RUNX1* KO selectively moved macrophages away from pro-inflammatory states (Mac5) and towards recovery-associated states (Mac4) (**Fig. 1I-J**). Given the dynamic transcriptional remodeling of macrophages observed in human cardiac recovery (**Fig. 1A-B**), we ranked all genes expressed in the macrophage compartment based on their correlation with ejection fraction. Genes were classified as negatively correlated if their expression increased as cardiac function declined and decreased as function improved; the opposite trend defined a positive correlation. Notably, *RUNX1* was among the most negatively correlated genes (**Fig. 1K**), showing elevated expression in heart failure (HF), reduced levels in patients who recovered cardiac function, and persistently high expression in non-recovering (NR) patients. These findings are consistent with our prior work showing that *RUNX1* expression dynamically changes in macrophages in human recovery validated with transcriptomics and RNA-scope in situ hybridization. These findings support the therapeutic hypothesis that targeting *RUNX1* within macrophages simultaneously impedes transcriptional networks associated with HF and contributes to the acquisition of recovery programs.

### *Runx1* deletion in macrophages leads to functional cardiac recovery

To selectively target *Runx1* in cardiac macrophages, we crossed *Runx1*^flox/flox^ mice to *Cx3cr1*^CreERT2^ mice, in which a tamoxifen-inducible Cre recombinase is driven by the *Cx3cr1* promoter. To assess the impact of *Runx1* deletion on the cardiac myeloid cell landscape in healthy hearts, we subjected control and *Runx1*^flox/flox^*Cx3cr1*^CreERT2^ animals to five days of continuous tamoxifen treatment. Flow cytometry analysis showed no differences in proportion of macrophages (CD64^+^), monocytes (Ly6c^low^ and Ly6c^high^), and neutrophils (Ly6g^+^) between control and *Runx1*^flox/flox^*Cx3cr1*^CreERT2^ animals (**Suppl. Fig. 2A**). Further gating within the mononuclear phagocyte compartment revealed no differences in CCR2^+^ (C-C chemokine receptor 2) and CCR2^-^ macrophages or dendritic cells (cDC1 and cDC2) between groups (**Suppl. Fig. 2A**).

To test the hypothesis that deleting *Runx1* in cardiac macrophages can facilitate recovery in a model of established heart failure, we utilized a closed-chest ischemia reperfusion injury (IRI) model^38^. Control and *Runx1*^flox/flox^*Cx3cr1*^CreERT2^ (*Runx1*-KO) animals were subjected to IRI (90 minutes of ischemia), followed by echocardiography at 3 weeks after IRI to establish the heart failure phenotype and quantify diminishment in LV ejection fraction, extent of LV dilation, and akinetic segment size (**Fig. 2A**). We then deleted *Runx1* in macrophages by administering tamoxifen beginning 3 weeks after IRI injury and assessed mice by echocardiography at 8 and 12 weeks post-IRI (**Fig. 2A**). Serial echocardiography showed that control mice displayed progressive LV remodeling with declining LV ejection fraction and increasing LV chamber dimensions. In contrast, *Runx1*-KO animals displayed remarkable and ongoing recovery of LV ejection fraction and reduced LV chamber dimensions at 8- and 12-weeks post-IRI (**Fig. 2B-D**). Trichrome staining in hearts 12-weeks post-IRI showed that that *Runx1*-KO animals had a significantly less collagen dense scar compared to controls (**Fig. 2E**).

**Figure 2.**
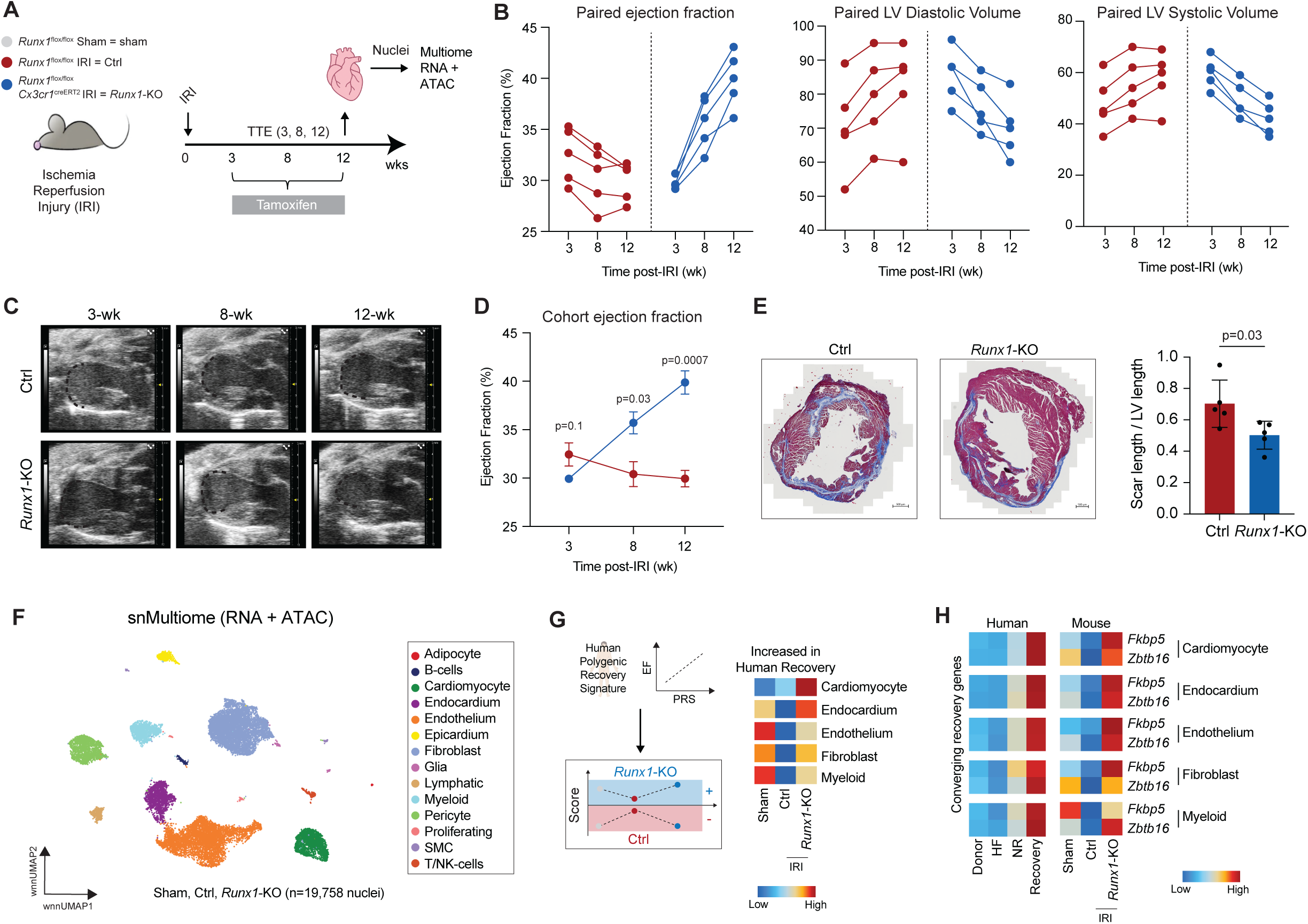
*Runx1* deletion in macrophages promotes functional and molecular recovery. (A) Study design showing *Runx1*^flox/flox^ *Cx3cr1*^CreERT2^ (*Runx1*-KO) and *Runx1*^flox/flox^ (WT) animals subjected to sham or ischemia reperfusion surgery with serial functional measurements with echocardiography at 3-, 8-, and 12-weeks post-IRI with continuous tamoxifen administration between 3-12 weeks. At 12 weeks post-IRI, sham, WT, and *Runx1*-KO animals were sacrificed and nuclei isolated for Multiome (RNA + ATAC) sequencing. (B) Paired EF, LV diastolic volume, and LV systolic volume measurements in WT and *Runx1*-KO animals post-IRI demonstrating significant functional recovery upon *Runx1* deletion. (C) Long axis paired still example images in Control and *Runx1*-KO animals post-IRI. (D) Grouped EF measurements in WT and *Runx1*-KO animals post-IRI demonstrating significant functional recovery upon *Runx1* deletion. (E) Quantification of myocardial scar burden/total LV circumference in *Runx1*-KO and WT animals 8-weeks post-IRI. (F) WNN UMAP embedding plot of sham, WT, and *Runx1*-KO animals colored by cell type. (G) Schematic depiction of a cell type specific human polygenic recovery score mapped onto mouse data (left). Heatmap of cell type specific positive PRS score from human recovery mapped onto mice grouped by sham, WT, and *Runx1*-KO animals (right). (H) *Fkbp5* and *Zbtb16* expression in human (left) and mouse recovery by cell type.

### *Runx1* deletion reprograms the cardiac transcriptional and epigenomic landscapes toward recovery

Given the functional and histological recovery seen in *Runx1*-KO animals, we next performed single nuclei multiome (RNA+ATAC) sequencing to characterize the transcriptional and epigenomic landscape (**Fig. 2A**). After quality control, peak calling (**Suppl. Fig. 3A-B**), and data normalization, we performed weighted nearest neighbor (WNN) clustering^46–51^ using gene expression and chromatin accessibility data (**Suppl. Fig. 3C**). Differential gene expression analysis identified fourteen major cell types based on canonical marker genes (**Fig. 2F**, **Suppl. Fig. 3C-G**). To determine whether *Runx1*-KO animals adopt a similar transcriptional signature to that seen in human recovery, we generated cell type specific polygenic recovery scores (PRS) based on genes positively correlated with LV ejection fraction in human hearts^6^ (**Fig. 2G**, **Suppl. Fig. 3H**). The PRS declined 3 weeks after reperfused MI in WT hearts, and conversely, increased across all major cell types (cardiomyocytes, fibroblasts, endothelium, endocardium, and myeloid cells) in *Runx1*-KO hearts highlighting cell autonomous and non-cell autonomous effects of macrophage *Runx1* deletion during recovery (**Fig. 2G**).

To systematically characterize regulatory dynamics in murine recovery we performed cell type specific differential gene expression, chromatin accessibility, and transcription factor (TF) motif analysis between control and *Runx1*-KO animals to investigate pathways implicated in recovery (**Suppl. Fig. 4A**). We found broad transcriptional and regulatory changes in cardiomyocyte, endocardium, endothelium, fibroblast, and myeloid cell populations (**Suppl. Fig. 4A**). Notably, ATAC-seq in *Runx1*-KO hearts showed similarities to healthy sham animals, but also, acquisition of a unique recovery state that differed from both sham and control IRI conditions (**Suppl. Fig. 4B**). To map regulatory changes to functional pathways, we used peak-to-gene linkage^49^ and performed gene ontology enrichment analysis on genes linked to peaks showing greater accessible in *Runx1*-KO hearts compared to control hearts. This enhancer-to-gene-to-pathway framework uncovered cell type–specific recovery programs (**Suppl. Fig. 4C**). Recovered cardiomyocytes showed enrichment in pathways related to cardiac muscle development, contractility, and hypertrophy; the endocardium in amide metabolism and histone modification; endothelium in regulation of cell migration, proliferation, and vasculogenesis; fibroblasts in environmental sensing, BMP signaling, and interferon alpha responsive genes; and macrophages in leukocyte activation, myeloid cell differentiation, and antigen presentation (**Suppl. Fig. 4B**). Finally, to explore conserved transcriptional signatures of recovery shared across cell types and between human and murine systems, we overlapped genes upregulated in *Runx1*-KO hearts with those enriched in human recovery. Among the shared genes, *FKBP5* and *ZBTB16* followed a conserved trajectory: modest expression in sham or non-failing donor hearts, increased expression in heart failure and control IRI hearts, and suppression in recovered and *Runx1*-KO hearts (**Fig. 2H**). Collectively, these data indicate that *Runx1*-KO hearts display a transcriptional and regulatory signature representative of human recovery.

### BRD4 controls *Runx1* expression and transcriptional network activity in macrophages

To further explore *RUNX1* regulation in a tractable murine model of cardiac recovery, we used a murine pressure overload model (TAC) in which heart failure can be pharmacologically reversed and re-induced. Mice subjected to transverse aortic constriction (TAC) were treated with JQ1, a small-molecule inhibitor that reversibly targets bromodomain and extraterminal domain (BET) proteins—key transcriptional regulators of pathological gene expression in the failing heart^31,52–54^. JQ1 treatment restored cardiac function to near-baseline levels, while withdrawal of the drug led to rapid functional decline^31,52,54^. In this model, *Runx1* expression was elevated in TAC, suppressed with JQ1 treatment, and reactivated upon JQ1 withdrawal^6,54^ suggesting a dependence on BET protein activity. To test the functional role of BET-dependent chromatin remodeling in regulating *Runx1* expression in cardiac macrophages, we genetically deleted the BET family member *Brd4* in macrophages. *Brd4*^flox/flox^ mice were crossed with *Cx3cr1*^CreERT2^ mice^31^ (**Fig. 3A**). We focused on BRD4 as it contributes to the transcriptional regulation of heart failure pathogenesis, including the control of pro-inflammatory gene expression in cardiac macrophages^31,52,53^. Mice were subjected to transverse aortic constriction (TAC), followed by one week of continuous tamoxifen treatment to delete *Brd4* in monocytes and macrophages^31^. On day 7 post-TAC—a timepoint at which the number of cardiac immune cells reaches its peak, CD45^+^ cells were isolated from sham, *Brd4*^flox/flox^, and *Brd4*^flox/flox^*Cx3cr1*^CreERT2^ hearts, and subjected to single-cell RNA sequencing (**Fig. 3A, Suppl. Fig. 5A,B**). *Runx1* expression in macrophages was elevated in TAC hearts compared to sham and was markedly reduced in *Brd4*-deficient hearts subjected to TAC (**Fig. 3B**). Interestingly, *Runx1* expression was selectively decreased in *Ccr2*^+^ macrophages and was not changed in *Ccr2*^-^ macrophages (**Fig. 3C, Suppl. Fig. 5C-E).** These findings suggest that BRD4-dependent regulation of *Runx1* is primarily restricted to monocyte-derived macrophages. To investigate the contribution of RUNX1 to the transcriptional and chromatin dynamics driven by BET proteins during heart failure and recovery, we first identified a set of 231 genes in macrophages that were upregulated in TAC, downregulated by JQ1 treatment, and re-upregulated upon JQ1 withdrawal. TF motif analysis of this gene set revealed significant enrichment of RUNX1 motifs among the top candidates (**Fig. 3D**). To complement this, we analyzed macrophage chromatin accessibility using single-cell ATAC-seq and identified distal regulatory regions with increased accessibility following JQ1 withdrawal (**Fig. 3E**), a condition that results in rapid decrease of cardiac function compared to continuous JQ1 treatment. These regions were also enriched for RUNX1 motifs, supporting a role for RUNX1 in chromatin reactivation during cardiac functional decline (**Fig. 3E**).

**Figure 3.**
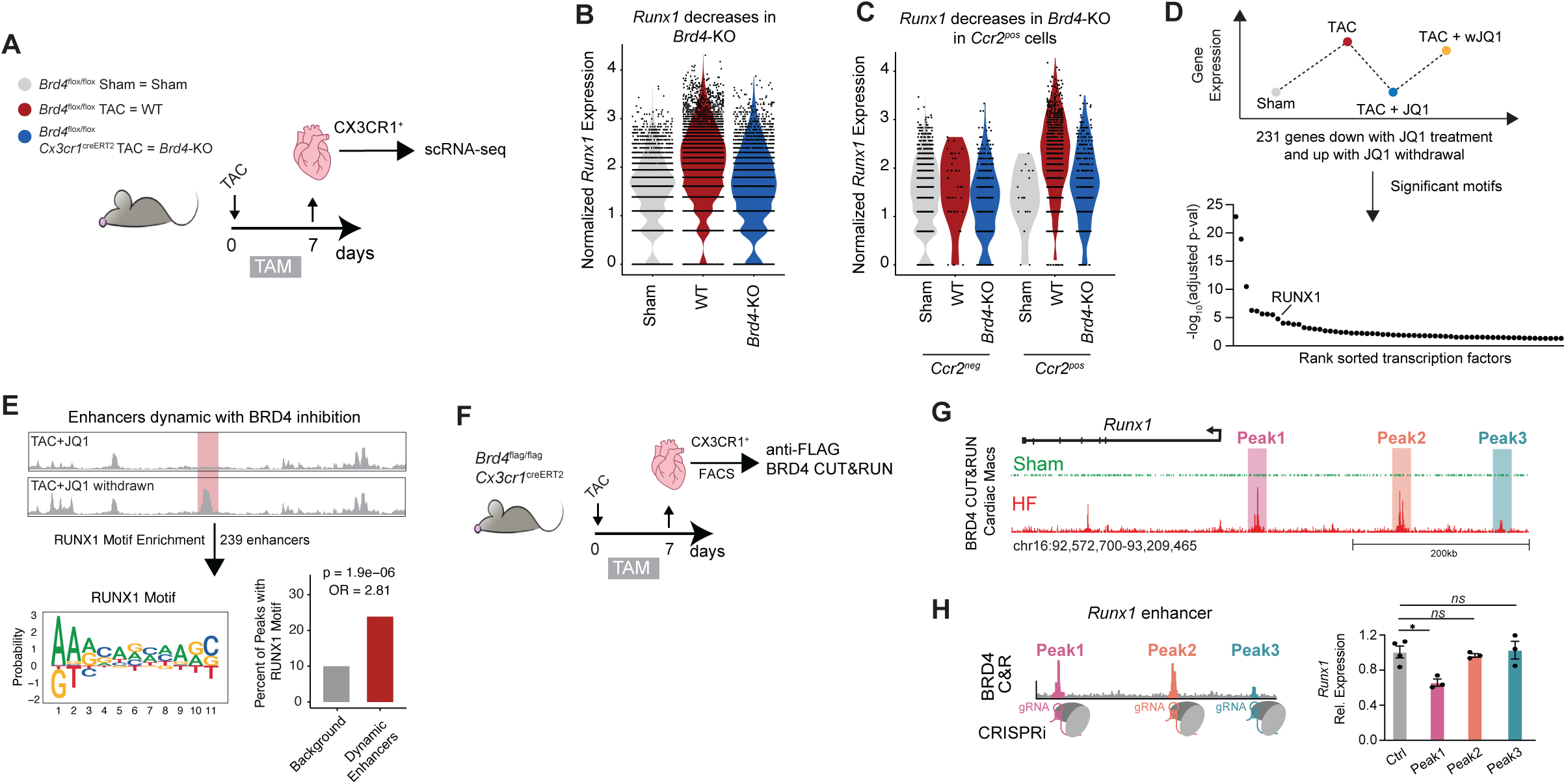
BRD4 controls stress-dependent *Runx1* transcription dynamically in recovery. (A) Study design for targeted deletion of *Brd4* in cardiac macrophages in a in vivo model of heart failure with isolation of macrophages for single cell RNA sequencing. (B) Violin plot for *Runx1* expression in macrophages from sham, *Brd4*^flox/flox^, and *Brd4*^flox/flox^ *Cx3cr1*^CreERT2^ animals. (C) Violin plot for *Runx1* expression grouped by *Ccr2*^pos^ and *Ccr2*^neg^ macrophages from sham, *Brd4*^flox/flox^, and *Brd4*^flox/flox^ *Cx3cr1*^CreERT2^ animals. (D) Schematic illustrating genes which are downregulated with JQ1 treatment and revert upon withdrawal of JQ1 (top) and transcription factor enrichment of dynamically regulated 231 genes. The adjusted p-value is computed using the Benjamini-Hochberg method for correction for multiple hypothesis testing. (E) Schematic showing distal regulatory regions with increased accessibility following JQ1 withdrawal (top) with motif enrichment showing the RUNX1 PWM and percent of peaks enriched with RUNX1 motifs relative to background (bottom). (F) Study design for in vivo BRD4 chromatin occupancy profiling with anti-FLAG BRD4 CUT&RUN from *Brd4*^flag/flag^ *Cx3cr1*^CreERT2^ animals post-TAC. (G) anti-FLAG BRD4 CUT&RUN identifying stress dependent recruitment to distal regulatory elements Peak1, 2, and 3 upstream of *Runx1*. (H) Series of CRISPRi perturbation experiments in RAW264.7 macrophages with a non-targeting control, guides targeting peaks1-3 with qPCR for *Runx1* expression.

To dissect the molecular mechanism by which BRD4 regulates *Runx1*, we used a knock-in mouse model in which a 3×FLAG epitope tag is fused to the N-terminus of *Brd4* (*Brd4*^flag/flag^)^31^. Heart failure was induced by TAC, and cardiac macrophages were sorted at day 7 post-surgery—a timepoint at which Brd4 deletion in macrophages led to reduced *Runx1* expression (**Fig. 3F-H**). We then performed anti-FLAG CUT&RUN (Cleavage Under Targets and Release Using Nuclease) to map Brd4 chromatin occupancy *in vivo* with high sensitivity and low input requirements (**Fig. 3F**). This analysis revealed three Brd4-enriched distal peaks within a large upstream region of the *Runx1* locus (**Fig. 3G**). To determine which of these regions functionally regulate *Runx1*, we leveraged CRISPR interference (CRISPRi) in a murine macrophage cell line (RAW 264.7) expressing dCas9-KRAB (**Suppl. Fig. 5G-I)**, enabling selective repression of individual cis-regulatory elements within the *Runx1* enhancer (**Fig. 3H**). Targeted repression of each individual candidate enhancer revealed that only inhibition of the Peak1 region significantly reduced *Runx1* expression, identifying this element as the Brd4-dependent cis-regulatory element controlling *Runx1* activation in macrophages (**Fig. 3H**).

### *Runx1* deletion in cardiac macrophages recapitulate a recovery phenotype

To characterize how *Runx1* deletion in macrophages shifts the immune landscape in our ischemic heart failure model, we sorted CD45^+^ cells from sham, control, and *Runx1*-KO hearts 8 weeks after IRI and performed single cell RNA-sequencing (**Fig. 4A**, **Suppl. Fig. 6A**). After quality control, clustering, and differential expression analysis we identified several immune cell subsets (**Fig. 4B**, **Suppl. Fig. 6B-C**). We then isolated the monocyte and macrophage populations for higher resolution cell state annotation and found 9 transcriptionally distinct cell states (**Fig. 4C**, **Suppl. Fig. 6D**). *Cx3cr1* expression confirmed all cell states were monocytes and macrophages, *Ccr2* expression was highest in the monocyte populations, and *Cd163* was enriched in Mac4-6 indicative of tissue resident macrophages enriched in sham animals^35^ (**Suppl. Fig. 6E**). Cell state composition showed that *Runx1*-KO hearts displayed marked expansion in Mono1 (enriched with type 1 interferon responsive genes) and a reduction in Mac2 (enriched with pro-inflammatory genes), which is highly similar to inflammatory macrophages (human Mac3, Mac5) reduced in human cardiac recovery (**Fig.1E**, **Fig. 4D-E**). Differential gene expression analysis between control and *Runx1*-KO monocytes and macrophages showed that *Runx1*-KO macrophages displayed increased expression of *Tsc22d3* and *Dusp1* (glucocorticoid signaling genes), *Cxcr4*, *H2-T23* (MHC-II gene), and the type 1 interferon responsive genes *Ifitm3, Ifitm6, Ifi27l2a, Ifi211, Ifi30, Ifi47, Irf7, Oasl2, Isg15, Oas1a,* and *Ly6e* (**Fig. 4F**). The above interferon-responsive genes were strongly enriched within Mono1, which expands in *Runx1*-KO hearts (**Suppl. Fig. 6F**). These data are consistent with our prior finding that type 1 interferon signaling in macrophages is protective following IRI^55^, highlighting the potential relevance of this population in cardiac recovery. Among the recovery associated genes, *Tsc22d3* showed the strongest enrichment in both human recovery and *Runx1*-KO hearts across all monocyte and macrophage cell states (**Fig. 4F, Suppl. Fig. 6G-H**). Similarly, *TSC22D3* expression was elevated in macrophages from patients who recovered (**Suppl. Fig. 6H**).

**Figure 4.**
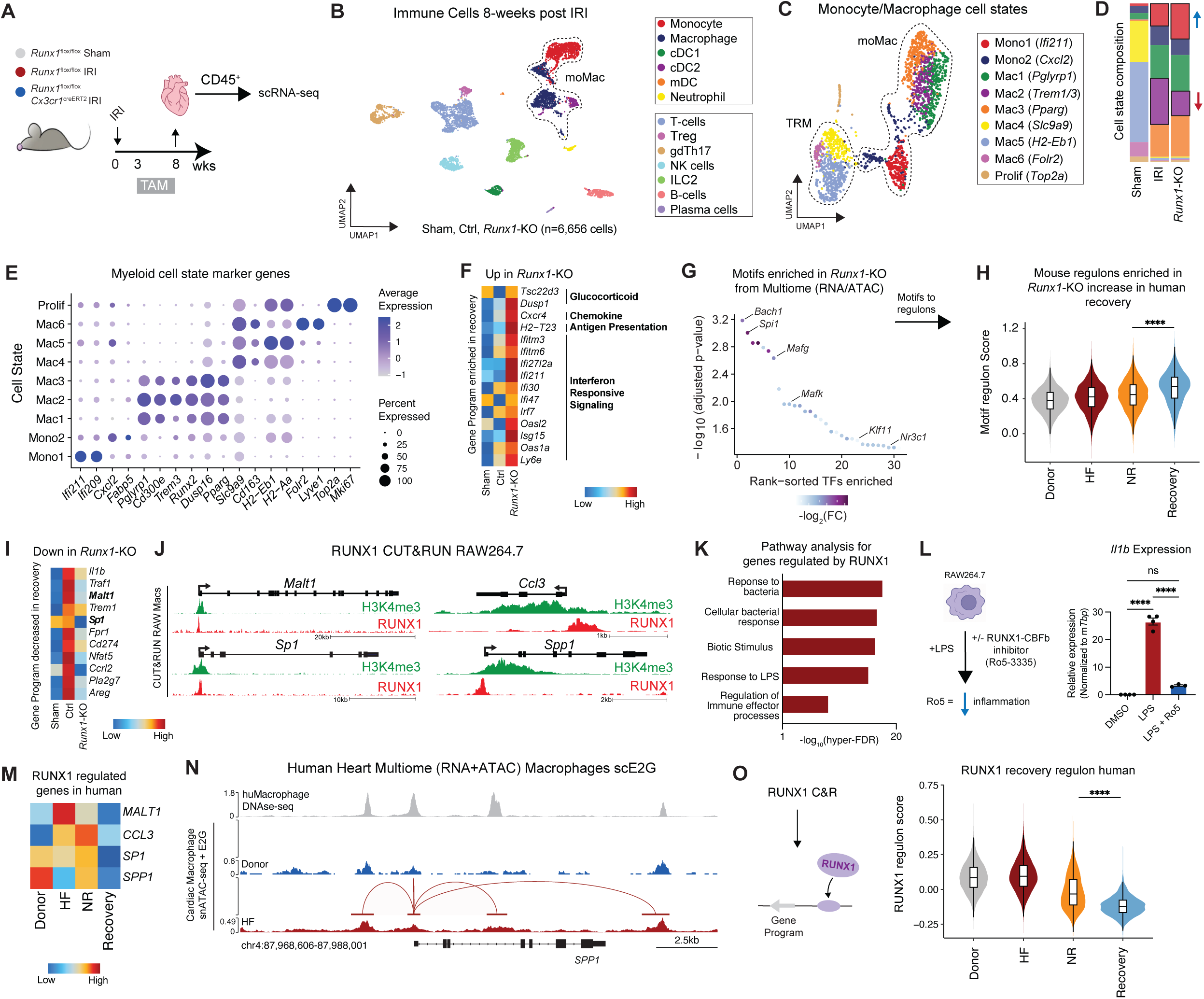
*Runx1* controls an inflammatory gene program and regulates macrophage state dynamics. (A) Workflow for isolation of immune cells for single cell gene expression profiling from WT and *Runx1*-KO animals 8-weeks post-IRI. (B) UMAP embedding plot of all CD45+ immune cells colored by cell type. (C) UMAP embedding plot for monocyte and macrophage cell state. Dotted lines outline monocyte derived and tissue resident macrophage populations. (D) Relative cell state composition in WT and *Runx1*-KO animals (right) colored by cell state. (E) Dot plot for top 2 marker genes for macrophage cell states. (F) Heatmap for genes upregulated in *Runx1*-KO relative to WT 8-weeks post-IRI grouped by functional significance. (G) TF motifs upregulated in *Runx1*-KO relative to WT 12-weeks post-IRI based on differential chromatin accessibility. (H) Gene set score for human ortholog genes within the motif regulon from (G). (I) Heatmap of genes downregulated in *Runx1*-KO relative to WT 8-weeks post-IRI. (J) Genomic tracts for RUNX1 and H3K4me3 CUT&RUN in RAW264.7 macrophages at key heart failure associated genes. (K) Gene Ontology pathway analysis for compendium of RUNX1 target genes obtained from anti-RUNX1 CUT&RUN in RAW264.7. (L) *Il1b* gene expression in untreated, LPS treated, and LPS + Ro5-3335 treated RAW264.7 macrophages. (M) Heatmap of example genes directly regulated by RUNX1 grouped by donor, heart failure, non-responder, and recovered human hearts. (N) Human macrophage DNAseq and scE2G links in Multiome (RNA+ATAC) data from human heart samples split by donor and HF at the *SPP1* locus showing a E2G link from a non-coding enhancer to the promoter. (O) Gene set score for human ortholog genes dynamic in recovery and under direct control of RUNX1 (from anti-RUNX1 CUT&RUN) decreases in human recovery.

To explore regulatory mechanisms underlying recovery associated genes in *Runx1*-KO macrophages, we performed transcription factor motif enrichment analysis in chromatin peaks that displayed increased accessibility in *Runx1*-KO macrophages. We found enrichment for Bach1, Spi1, Mafg, Mafk, Klf11, and Nr3c1 motifs (**Fig. 4G**). To assess whether these TFs might also be implicated in macrophage transitions seen in human cardiac recovery, we used these motifs to construct a macrophage recovery regulon score based on human orthologs. This analysis indicated significant enrichment in macrophages from patients who recovered compared to those with heart failure (**Fig. 4H**). Taken together, these findings indicate that *Runx1* deletion in cardiac macrophages recapitulates human recovery signatures.

We next investigated how *Runx1* deletion reshapes inflammatory macrophages subsets. The reduction in Mac2 abundance in *Runx1*-KO hearts (**Fig. 4D**) suggested a shift away from inflammatory macrophage states. To better understand the origin of Mac2, we leveraged published monocyte lineage tracing data (*Ccr2*^CreERT2^*Rosa26*^tdTomato^) using our IRI model^55,56^, and found that Mono1-2 and Mac1-3 mapped to monocyte-derived populations (**Suppl. Fig. 7A-D**). Differential expression analysis revealed that *Runx1* deletion in macrophages led to downregulation of pro-inflammatory genes including *Il1b, Traf1, Malt1, Trem1, Sp1, Fpr1, Cd274, Nfat5, Ccrl2, Pla2g7,* and *Areg* relative to control hearts post-IRI. Each of these genes were markedly increased in macrophages from control hearts following IRI compared to sham hearts (**Fig. 4I**).

To identify genes under direct control of RUNX1, we performed anti-RUNX1 CUT&RUN in RAW264.7 macrophages to map RUNX1 chromatin occupancy (**Fig. 4J** and **Suppl. Fig. 7E**). Notably, we found evidence of direct RUNX1 binding at the promoters (confirmed with H3K4me3 marks) of genes such as *Malt1* and *Sp1*, which were also dynamically regulated *in vivo* upon *Runx1* deletion (**Fig. 4J**). We further identified RUNX1 binding at promoters and enhancers of *Ccl3* and *Spp1*, mediators implicated in adverse remodeling and previously associated with poor cardiac recovery in humans^6^ (**Fig. 4J**). Using monocyte lineage tracing data, we confirm that *Ccl3* and *Spp1* were enriched in *Ccr2*^+^ monocyte-derived macrophages in mice and corresponding populations in humans^25,55^ (**Suppl. Fig. 7B-D**). In human MI and heart failure, *CCL3* and *SPP1* expression is elevated in macrophages expressing *TREM1* which are similar to murine Mac2, which are both diminished in recovery (**Fig. 4D**, **Suppl. Fig. 7C-D**). Notably, we previously demonstrated that TREM1 signaling drives CCL3 expression, amplifying inflammatory responses during IRI^57^. To characterize the broader implications of RUNX1 regulation within macrophages we extracted all genes under direct control of RUNX1 and performed gene ontology analysis and found enriched response to bacteria, LPS, and effector immune processes (**Fig. 4K**, **Suppl. Fig. 7E**). We next tested whether this inflammatory program is pharmacologically targetable. Using Ro5-3335, a small molecule inhibitor that blocks RUNX1–CBF complex formation^39^, we treated RAW264.7 macrophages with LPS and either DMSO or Ro5-3335. This treatment led to a dose-dependent reduction in IL-1β and IL-6 at both the RNA and protein levels (**Fig. 4L, Suppl. Fig. 7F**), demonstrating effective suppression of inflammatory gene expression.

Importantly, several direct RUNX1 targets, including *MALT1, CCL1, SP1,* and *SPP1*, which are downregulated in *Runx1*-KO animals, were also suppressed in human recovery^6^ (**Fig. 4M**). To assess disease-specific chromatin regulation at these RUNX1 targets in humans we used single-cell macrophage scE2G maps from non-failing donors and heart failure patients to identify disease specific E2G links at the *SPP1* locus^25,43^ (**Fig. 4N**). Finally, we used our RUNX1 CUT&RUN data along with dynamic human recovery gene expression to construct a RUNX1 regulon, comprising direct targets that are transcriptionally modulated during recovery. We found a significant reduction in regulon activity in patients who recovered, compared to those with persistent heart failure (**Fig. 4O**). Collectively, these data suggest that RUNX1 controls recovery and inflammatory heart failure modules in cardiac macrophages that are conserved between humans and mice.

### *Runx1* deletion in macrophages remodels cardiomyocytes

To assess non-cell autonomous effects of macrophage *Runx1* deletion on cardiomyocytes, we isolated cardiomyocyte nuclei from sham hearts, control hearts 12 weeks post-IR, and *Runx1*-KO hearts 12 weeks post-IRI and performed full-length transcriptome profiling using the ICELL8 platform^60^ (**Fig. 5A-B**, **Suppl. Fig. 8A**). Following quality control, clustering, and differential expression analysis, we annotated the major cell types (**Fig. 5B**, **Suppl. Fig. 8A-C**). Notably, we detected more unique genes per nuclei with greater sensitivity as compared to standard single nuclei RNA sequencing workflows (**Suppl. Fig. 8D-F**). We performed differential expression analysis between control and *Runx1*-KO conditions and uncovered a transcriptional shift in the *Runx1*-KO group suggestive of adaptive remodeling (**Fig. 5B-D**). Cardiomyocytes from *Runx1*-KO hearts were enriched in pathways involved in enhanced cardiac contraction and calcium cycling while pathways linked to oxidative phosphorylation were decreased (**Fig. 5C**). Notably, cardiomyocytes from *Runx1*-KO hearts upregulated genes critical for calcium handling (*Cacna1c, Cacnb2, Ryr2, Camk2d,* and *Slc8a1*)^61^, sarcomere structure and contractility (*Tnnt2, Ttn, Myh6,* and *Myh7*)^62^, and cytoskeletal organization (*Sorbs1, Dmd, Ldb3*)^63,64^. In parallel, we observed upregulation of transcriptional and splicing regulators such as *Rbm20*, *Rbm24*, and *Myocd*^65–67^, as well as the lncRNAs *Malat1* and *Mhrt*. Cardiomyocytes from *Runx1* KO hearts also exhibited restoration of *Myh6*, *Myh7*, *Ttn, Tnnt2*, and *Mir133a* expression. Conversely, genes involved in mitochondrial oxidative phosphorylation and ATP synthesis^68^ including *mt-Nd1–6, Cox4i1, Atp5b*, and *Ndufa4*, were increased in control cardiomyocytes and reduced to sham levels in cardiomyocytes from *Runx1*-KO hearts. possibly suggesting a shift away from oxidative metabolism (**Fig. 5D**).

**Figure 5.**
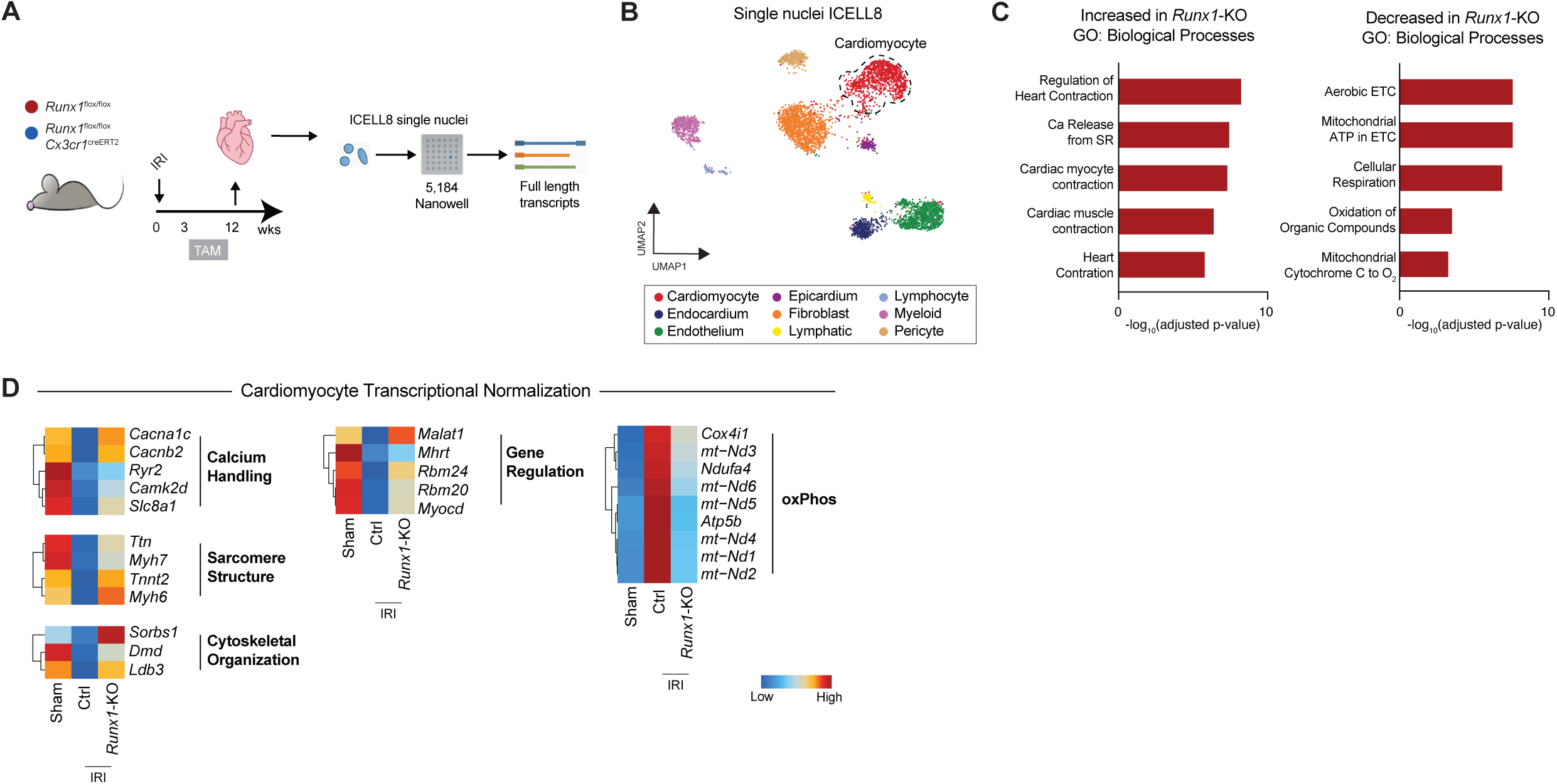
Recovered macrophages promote adaptive remodeling in cardiomyocytes. (A) Study design for nuclei isolation for ICELL8 single nuclei RNA-seq in nanowells for deep transcriptional profiling in WT and *Runx1*-KO animals 12-weeks post-IRI. (B) UMAP embedding plot of sham, from WT and *Runx1*-KO animals 12-weeks post-IRI colored by cell type. (C) Gene ontology biological pathway analysis for cardiomyocytes using differentially expressed in WT and *Runx1*-KO animals 12-weeks post-IRI. (D) Heatmap of differentially expressed genes between WT and *Runx1*-KO animals grouped by functional relevance. EF = ejection fraction. IRI = ischemia reperfusion injury. LV = left ventricle. WNN = weighted nearest neighbor. TAM = tamoxifen.

### Deletion of *Runx1* in macrophages rewires fibroblast states

We next aimed to prioritize the cell types in *Runx1*-KO hearts that most closely resemble the human recovery state. To do this, we computed the cosine similarity of transcriptional profiles between each cell type in *Runx1*-KO hearts and recovered human hearts (**Fig. 6A**). We found that macrophages and fibroblasts from *Runx1*-KO hearts show the strongest transcriptional similarity to human recovery (**Fig. 6A**). To more deeply characterize shifts in fibroblast cell states, we sorted fibroblasts from sham hearts and control and *Runx1*-KO hearts 8 weeks post-IRI and performed single cell RNA-sequencing (**Fig. 6B**, **Suppl. Fig. 9A**). After quality control, clustering, and differential expression analysis we identified 10 transcriptionally distinct fibroblast cell states (**Fig. 6C**, **Suppl. Fig. 9B-C**). Cell state composition analysis revealed that *Runx1*-KO hearts contained a relative reduction in Fib1 and Fib2, which were marked by high expression of *Postn* and *Comp* (classical markers of fibroblast activation)^69–71^, respectively (**Fig. 6C**). Next, we constructed a fibrosis gene signature (*Postn, Meox1, Runx1, Col1a1, Comp, Aebp1, Acta2, Thbs4, Cthrc1, Acta2,* and *Ltbp2*) using genes dysregulated in fibrotic tissues^25,31,54,69–75^ and found strongest enrichment in Fib1 and Fib2 (**Fig. 6D**). These activated fibroblast related genes were absent in sham fibroblasts, increased in control hearts post-IRI, and reduced in *Runx1*-KO hearts post-IRI (**Fig. 6E**). Pathway analysis of genes downregulated in Fib1 and Fib2 because of macrophage *Runx1* deletion showed enrichment for extracellular matrix and TGF-β signaling (**Fig. 6F**). Similarly, an activated fibroblast signature derived from human heart failure (*POSTN, COMP, FAP, COL1A1, THBS4, COL3A1*)^25^ localized to Fib1 and Fib2 (**Suppl. Fig. 9D**). Among these genes, *Fap* expression was significantly reduced in fibroblasts from *Runx1*-KO hearts, indicating suppression of pro-fibrotic fibroblast activation^25,76,77^ (**Suppl. Fig. 9E**).

**Figure 6.**
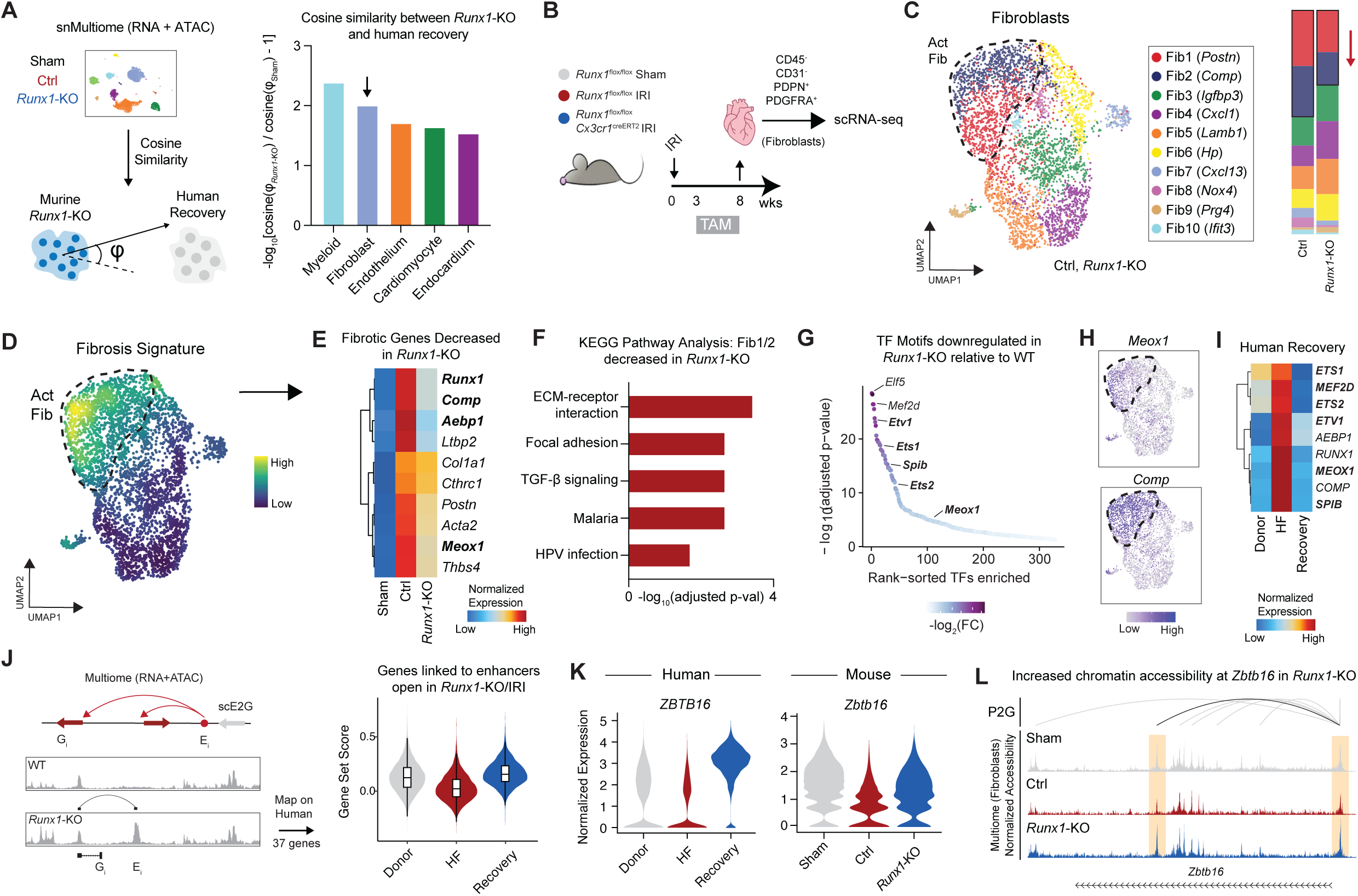
*Runx1* deficient animals suppress pro-fibrotic gene programs. (A) Schematic of cosine similarity computation between *Runx1*-KO animals post-IRI and human recovery (left) and quantification of a cosine similarity score by cell type (right). A higher value indicates greater similarity between recovered *Runx1*-KO animals and recovered human hearts. (B) Workflow for isolation of cardiac fibroblasts for single cell gene expression profiling from WT and *Runx1*-KO animals 8-weeks post-IRI. (C) UMAP embedding plot of fibroblasts colored by cell state (left) and relative cell state composition in WT and *Runx1*-KO animals (right). (D) Fibrosis gene set (*Postn, Meox1, Runx1, Col1a1, Comp, Aebp1, Acta2, Thbs4, Cthrc1, Acta2,* and *Ltbp2*) score density plot on fibroblast UMAP. (E) Heatmap of fibrosis genes downregulated in murine recovery grouped by sham, WT, and *Runx1*-KO. (F) KEGG pathway analysis for Fib1/2 marker genes downregulated in *Runx1*-KO. The adjusted p-value is computed using the Benjamini-Hochberg method for correction for multiple hypothesis testing. (G) TF motifs downregulated in *Runx1*-KO relative to WT animals 12-weeks post-IRI based on differential chromatin accessibility. (H) *Meox1* and *Comp1* expression restricted to activated Fib1/2 states. (I) Heatmap of pro-fibrotic genes and transcription factors decreased in human recovery. (J) Schematic of scE2G map in macrophages from WT and *Runx1*-KO animals 12-weeks post-IRI identifying genes links to enhancers more accessible in *Runx1*-KO relative to WT (left) and gene set score for human ortholog showing increased expression in human recovery (right). (K) Violin plot of *Zbtb16* expression in human and mouse recovery. (L) Chromatin accessibility and peak-gene linkage at the *Zbtb16* locus in sham, WT, and *Runx1*-KO fibroblasts; highlighted segment shows an intronic regulatory element more accessible in sham and *Runx1*-KO fibroblasts linked to the *Zbtb16* promoter. IRI = ischemia-reperfusion injury. TAM = tamoxifen. Act Fib = activated fibroblasts. scE2G = single cell enhancer gene. HF = heart failure.

To explore upstream regulators of fibroblast state changes, we performed transcription factor motif enrichment analysis in chromatin peaks downregulated in *Runx1*-KO relative to control hearts post-IRI and found enriched *Elf5*, *Mef2d*, *Ets1*, *Spib*, *Ets2*, and *Meox1* (**Fig. 6G**). Among these genes, *Meox1* was previously implicated in fibroblast activation^31,54^ and localized to the Fib1 and Fib2 cluster overlapping with *Comp* expression (**Fig. 6H**). We next asked whether a human pro-fibrotic program containing an overlapping set of genes (*ETS1*, *MEF2D*, *ETS2*, *ETV1*, *AEBP1*, *RUNX1*, *MEOX1*, *COMP*, and *SPIB*) displayed similar dynamic regulation in human cardiac recovery. These genes were absent in non-failing hearts, upregulated during heart failure, and decreased again in recovery, mirroring the murine transition (**Fig. 6I**).

To uncover the regulatory elements positively associated with recovery, we performed differential chromatin accessibility analysis between fibroblasts from control and *Runx1*-KO hearts to prioritize a set of putative enhancers that become more accessible in recovered fibroblasts. Using peak-to-gene linkage, we associated these regions with nearby genes and derived a recovery gene signature (**Fig. 6J**). Human orthologs of this signature were significantly increased in fibroblasts from human recovery relative to failing hearts (**Fig. 6J**). Gene ontology analysis showed enrichment of metabolic pathways, cell differentiation, and response to external stimuli (**Suppl. Fig. 9F**). Among the recovery associated genes, *Zbtb16* showed the strongest enrichment in human recovery and *Runx1*-KO hearts across all fibroblast cell states (**Fig. 6K, Suppl. Fig. 9G**). Multiome (RNA/ATAC) showed dynamic chromatin accessibility around the *Zbtb16* locus. Specifically, there is high chromatin accessibility within an intronic regulatory element linked to the promoter in sham fibroblasts, which diminished in control fibroblasts post-IRI, and increased in fibroblasts from recovered *Runx1*-KO hearts (**Fig. 6L**). Collectively, these findings demonstrate that *Runx1* deletion in macrophages imposes robust effects on cardiac fibroblast cell state acquisition and gene regulatory networks that mirrors what is observed in human cardiac recovery.

### Runx1 small molecule inhibition facilitates cardiac recovery in vivo

Finally, to the test the therapeutic potential of targeting *Runx1* in chronic heart failure, we utilized Ro5-3335, a small molecule inhibitor that blocks RUNX1–CBFb complex formation^39^. WT animals were subjected to IRI (90 minutes of ischemia), followed by echocardiography at 3 weeks after IRI to establish the heart failure phenotype and quantify diminishment in LV ejection fraction, extent of LV dilation, and akinetic segment size (**Fig. 7A-C**). We then treated animals with either control or Ro5-3335 subcutaneously beginning 3 weeks after IRI and assessed cardiac function by echocardiography at 12 weeks post-IRI (**Fig. 7A-E**). Serial echocardiography showed that untreated mice displayed progressive LV remodeling with declining LV ejection fraction and increasing LV chamber dimensions. In contrast, Ro5-3335 treated animals displayed remarkable and ongoing recovery of LV ejection fraction, reduced LV chamber dimensions, and a smaller akinetic area at 12 weeks post-IRI (**Fig. 7B-E**). To assess changes in infarct size and fibrosis, we performed trichrome staining and found that that Ro5-3335 treated animals had significantly less collagen dense scar compared to untreated animals (**Fig. 7F**).

**Figure 7.**
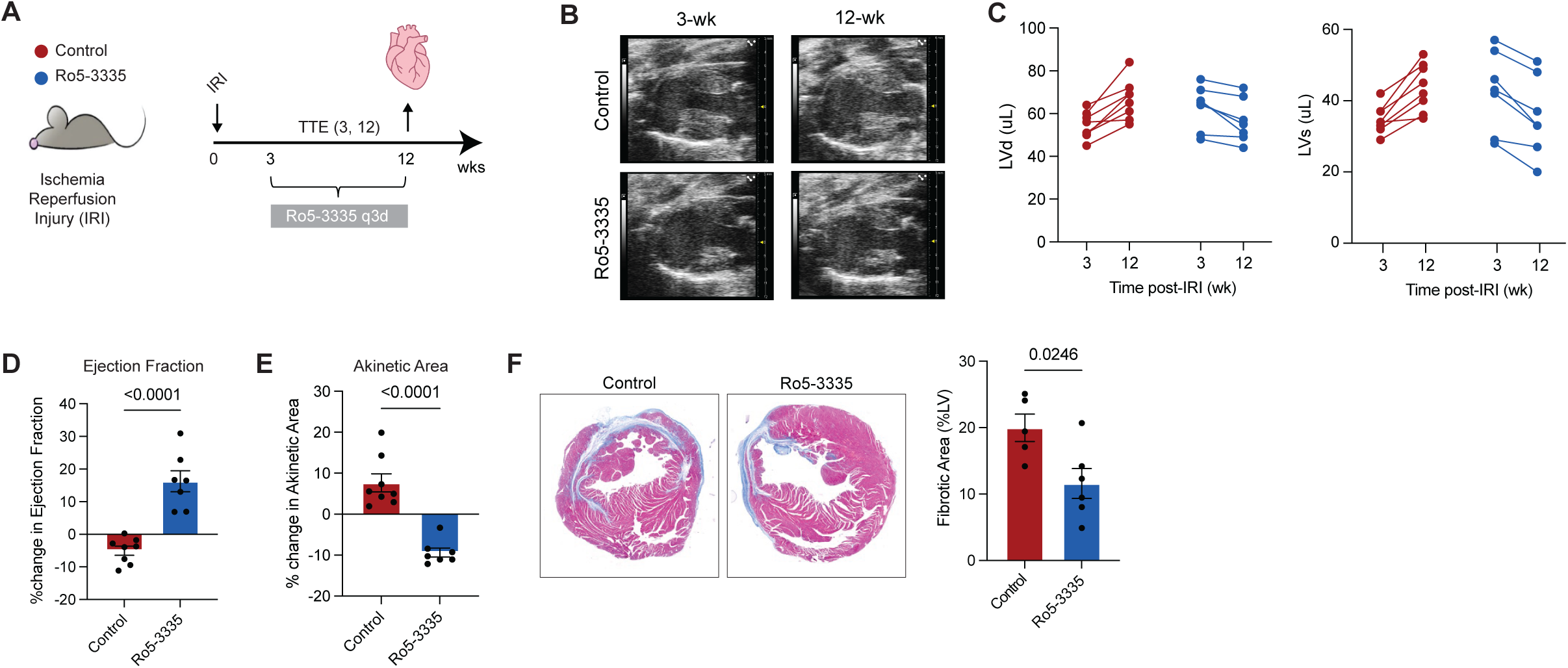
Small molecule Runx1 inhibition in vivo facilitates cardiac recovery. (A) Study design showing untreated and Ro5-3335 treated animals subjected to sham or ischemia reperfusion surgery with serial subcutaneous Ro5-3335 administration between 3-12 weeks. (B) Long axis paired still example images in untreated and Ro5-3335 treated animals post-IRI. (C) Paired LV diastolic and LV systolic volume measurements in untreated and Ro5-3335 treated animals post-IRI demonstrating significant functional recovery in the treatment group. Paired percent change in (D) ejection fraction and (E) akinetic area from 3 to 12 weeks post-IRI in untreated and Ro5-3335 treated animals. (F) Quantification of myocardial fibrotic area in untreated and Ro5-3335 treated animals 12-weeks post-IRI.

## Discussion

### Organ recovery as a transcriptional state of adaptation

Recovery of chronically diseased organs represent a clinical important but poorly understood biological process. While transcriptional pathways that drive organ failure are increasingly studied, the regulatory programs that enable reversal of organ dysfunction remain largely undefined. Heart failure provides a unique clinical setting to study organ recovery, as some patients supported with LVADs experience sustained restoration of cardiac function^1–4,6–11,16,17^. The availability of heart tissue in this setting offers an opportunity to define the molecular mechanisms that reverse organ dysfunction in humans. Elucidating how cells reprogram toward a reparative state may unlock new strategies to restore function in damaged tissues.

Through integrative multiomic profiling and network-based prioritization of healthy, failing, and recovered human hearts, we discover *RUNX1* within macrophages as a putative regulator of myocardial recovery. In vivo, we show that targeted deletion of *Runx1* in cardiac macrophages leads to functional, physiological, and molecular recovery with enrichment of human recovery gene programs across multiple cardiac cell types. We use single cell gene expression and chromatin profiling, in vivo TF occupancy assays, and a series of CRISPRi perturbations to mechanistically identify BRD4 as an upstream regulator of *Runx1* in cardiac recovery. Finally, we show that RUNX1 governs a druggable gene regulatory network in macrophages that impedes cardiac recovery by maintaining macrophages in an inflammatory state that imposes robust effects across major cardiac cell populations.

### Macrophage-specific *Runx1* deletion facilitates functional and molecular recovery

Previous work has demonstrated that modulating innate immune activation can reduce adverse remodeling in models of acute cardiac injury^25,31,34,35,55,56,78–81^. These studies have established that inhibiting nodal pathways in monocytes and macrophages can suppress pathological cell state transitions in stromal populations like cardiomyocytes and fibroblasts to reduce fibrosis and prevent organ dysfunction. However, whether the immune system can be harnessed to actively promote functional recovery in the failing heart remains unclear. Here, we used a chronic ischemic heart failure model initiated by IRI^38^ and showed that targeted deletion of *Runx1* in cardiac macrophages after the establishment of heart failure facilitates functional and structural recovery of the heart. Single-cell transcriptomic and epigenomic profiling revealed broad reprogramming across cardiomyocytes, fibroblasts, endothelial cells, and immune cells. We found that *Runx1*-macrophage deficient hearts adopt a transcriptional signature remarkably similar to that observed in human myocardial recovery. This cross-species convergence reinforces the concept that recovery is not a simple reversion to the healthy state, but rather an evolutionarily conserved adaptive program that emerges under conditions of stress reversal and functional improvement.

### BRD4 controls stress-inducible RUNX1 expression in monocyte-derived macrophages

BET bromodomain proteins have emerged as epigenetic regulators of inflammation in the heart and other tissues. Prior studies have shown that BET inhibition with the small-molecule JQ1 can reshape chromatin landscapes and suppress pathological gene expression in heart failure^31,52,54^. Furthermore, recent work has shown that BRD4 regulates macrophage plasticity in the failing heart by directly binding to key regulatory elements to facilitate pathological gene transcription^31^. Here, we show that *Runx1* expression is dynamically regulated by BRD4 in both human and murine recovery and that *Brd4* deletion in *Cx3cr1*+ macrophages selectively suppresses *Runx1* expression in monocyte-derived, but not tissue-resident macrophages. Furthermore, we use in vivo BRD4 occupancy mapping to show stress dependent recruitment at multiple upstream regulatory elements at the *Runx1* locus. Using a series of CRISPRi perturbations we discovered the precise regulatory element through which BRD4 controls *RUNX1* transcription. Together, these findings define a BRD4–RUNX1 regulatory circuit that links inflammatory stress to the maintenance of maladaptive macrophage states.

### RUNX1 controls a conserved inflammatory regulon in cardiac macrophages

Macrophages are uniquely positioned to integrate environmental cues and orchestrate tissue responses to injury. Our work demonstrates that in both human recovery and murine *Runx1* deletion models, suppression of a RUNX1-driven macrophage program is a hallmark of the recovery in both humans and mice. RUNX1 was highly expressed in failing hearts and reduced in recovered states. CUT&RUN mapping revealed direct binding of RUNX1 at promoters of key pro-inflammatory genes—including *Malt1, Sp1, Ccl3*, and *Spp1*—all of which were dynamically regulated upon *Runx1* deletion and enriched in CCR2⁺ TREM1⁺ macrophages associated with adverse outcomes in heart failure^35,57,82,83^. Conversely, macrophage-specific deletion of *Runx1* resulted in the expansion of glucocorticoid-responsive and interferon-enriched macrophage states, mirroring cell states found in human recovery. Indeed, we have previously shown that type 1 interferon responsive macrophages are cardioprotective^55^, underscoring the importance of this population in recovery. By integrating in vivo *Runx1* deletion, RUNX1 chromatin occupancy experiments, and enhancer-gene mapping, we constructed a regulon of RUNX1 target genes and show that the expression of these genes is markedly diminished in recovered human hearts. These results indicate that RUNX1 defines a disease-associated regulon that governs macrophage state transitions during recovery.

### Recovered macrophages impart adaptive changes in cardiomyocytes to promote recovery

Cardiac function is driven by an admixture of cell state diversity, ECM composition, and cellular synchronization converging on the contractile unit of the heart, the cardiomyocytes. Prior studies have used proteomics and ex vivo physiology assays to characterize myocyte biology in failing, mechanically unloaded, and recovered human hearts^20,84–88^. Deep transcriptomic profiling revealed that recovered cardiomyocytes upregulated genes critical for calcium handling, contractility and sarcomere structure, cytoskeletal organization, and partial reactivation of fetal-like programs with a concomitant shift away from oxidative metabolism. Reduced oxidative phosphorylation in recovered hearts may reflect an adaptive metabolic shift that enhances energy efficiency under stress, consistent with studies showing that failing and mechanically unloaded hearts often downregulate mitochondrial metabolism as a compensatory response to limit reactive oxygen species production and preserve cellular integrity^89–92^. Additionally, prior work has shown that modulation of mitochondrial biogenesis and oxidative phosphorylation via PGC-1α and ERR signaling plays a central role in maintaining myocardial energy homeostasis, and that partial downregulation of these pathways during recovery may protect against energetic overload and maladaptive remodeling^93^. Together, these findings suggest that macrophage-specific *Runx1* deletion promotes a cardiomyocyte phenotype optimized for energy efficiency and resilience.

### Recovered macrophages suppress pro-fibrotic programs in fibroblasts via cell–non-autonomous mechanisms

Previously we and others have shown that macrophages regulate fibroblast cell state transitions through cell non-autonomous signaling via secretion of pro-inflammatory and fibrotic chemokines and cytokines such as IL-1β^25,31^. In addition to macrophages, fibroblasts harbor the greatest transcriptional changes in heart failure and myocardial recovery. We discovered that during murine recovery, *Runx1-*macrophage deficient animals downregulate pro-fibrotic gene programs in activated fibroblasts through chromatin remodeling and reduced accessibility of TF motifs predicted to bind *MEOX1* and *RUNX1*, genes which have been previously implicated in cardiac fibroblast activation and fibrosis^6,25,26,31,54^. Furthermore, we find that recovered animals have reduced *Fap* expression which has been causally linked to chronic fibrosis and organ dysfunction^25,76,77,94^. Notably, recovered fibroblasts showed upregulation of gene programs seen in human recovery at the RNA and chromatin level – specifically, *Runx1*-macrophage deficient animals had enriched chromatin signal at a regulatory element linked *Zbtb16* not present in failing hearts. Prior studies have implicated *Zbtb16* as a transcriptional regulator of fibroblast quiescence and stress adaptation^95,96^. These results support a model in which macrophage reprogramming can rewire fibroblast transcriptional identity to promote a reparative state.

### Therapeutic implications and translational potential

Our findings reveal that targeting RUNX1 in cardiac macrophages is sufficient to trigger transcriptional and functional features of myocardial recovery. RUNX1 forms a transcriptional complex with CBFβ to bind regulatory elements and promoters and drive gene transcription^97^. Ro5-3335, a small molecule inhibitor, disrupts binding of RUNX1 with CBFβ thereby preventing downstream gene transcription^39^. Through in vivo genetic manipulation and chromatin mapping we show that *RUNX1* regulates a broad catalogue of pro-inflammatory gene programs, and treatment with Ro5-3335 leads to a dose dependent reduction in RNA and protein levels of canonical pro-inflammatory cytokines in stressed macrophages. Finally, we show that in vivo inhibition of the CBFβ-RUNX1 complex with a small molecule facilitates cardiac recovery in vivo. The selectivity of RUNX1 regulation in monocyte-derived macrophages offers a cell-state–specific opportunity to attenuate pathogenic inflammation without broadly suppressing macrophage function.

### Limitations of the study

Our study is not without limitations. First, while we show that *Runx1* deletion in *Cx3cr1*+ macrophages promotes cardiac recovery, this lineage includes monocyte-derived and some tissue-resident macrophages, and we cannot pinpoint the exact macrophage population where RUNX1 exerts its effects. Future studies using dual recombinase or temporally restricted Cre systems will help refine the ontogeny-specific contributions of RUNX1. Second, while we identify RUNX1 as a network-correcting node using in silico and in vivo perturbations, we have not evaluated the long-term safety or off-target effects of RUNX1 inhibition in chronic disease models or aged animals. Given the broad role of RUNX1 in hematopoiesis and immunity, further evaluation of toxicity and systemic immune modulation is warranted. Finally, although we identify conserved recovery signatures between human and murine hearts, human samples were limited in number and additional work in diverse patient populations and recovery settings will be necessary to validate generalizability of these findings.

### Conclusions

Collectively, these studies implicate inhibition of RUNX1 activity as a therapeutic target to facilitate cardiac recovery and uncover a mechanism whereby modulation of macrophage gene regulatory programs orchestrates recovery of heart function through suppression of inflammation, diminished fibroblast activation, and cardiomyocyte adaptation.

## Acknowledgments

KL is supported by the Washington University in St. Louis Rheumatic Diseases Research Resource-Based Center grant (NIH P30AR073752), the National Institutes of Health [R01 HL138466, R01 HL139714, R01 HL151078, R01 HL161185, R35 HL161185], Leducq Foundation Network (#20CVD02), Burroughs Welcome Fund (1014782), and Children’s Discovery Institute of Washington University and St. Louis Children’s Hospital (CH-II-2015-462, CH-II-2017-628, PM-LI-2019-829), Foundation of Barnes-Jewish Hospital (8038-88), and generous gifts from Washington University School of Medicine. JMA is supported by the Washington University School of Medicine Medical Scientist Training Program and Leducq Foundation Network Seed Grant (#20CVD02). M.A. was supported by the American Heart Association Second Century Award (24SCEFIA1252167), the Longevity Impetus Grant, and he is the Dario and Irina Sattui Investigator at the Gladstone Institutes. MUS acknowledges the support of an NSF Graduate Research Fellowship (DGE-1656518) and a graduate fellowship award from Knight-Hennessy Scholars at Stanford University. JME acknowledges support from NHLBI R01HL159176 and the Novo Nordisk Foundation Center for Genomic Mechanisms of Disease (NNF21SA0072102). Study design schematics were created in BioRender.com and using ChatGPT. The study was partially sponsored by a sponsored research agreement from Amgen. We thank the Genome Technology Access Center at the McDonnell Genome Institute at Washington University School of Medicine for help with genomic analysis. The Center is partially supported by NCI Cancer Center Support Grant #P30 CA91842 to the Siteman Cancer Center. This publication is solely the responsibility of the authors and does not necessarily represent the official view of NCRR or NIH.

## Author Contributions

JA and KL conceived the study. JA, MA, and KL drafted the manuscript. JA, MUS, TY, CJML, VX, and MA performed computational analyses. YA, AZ, CP, CYL, and MA performed all experiments with the BRD4 mice. JA, YLP, DP, JB, FS, NH, LL, TH, AP, JC, US, and IR performed in vivo studies. YA, AZ, MA, and JA performed CRISPR experiments. JA, KRK, and AB performed ICELL8 and Multiome experiments. LB assisted with animal breeding. MHG and AD performed ex vivo cardiomyocyte experiments. JN, CW, and AK performed animal surgeries and echocardiography. JC, CW, CML, DK, DLM, JN, IR, SJ, BA, JME, RF, RS, RSYF, SGD, and FL aided in data interpretation. All authors contributed to the experimental design, data analysis and interpretation as well as manuscript production. KL is responsible for all aspects of this manuscript including experimental design, data analysis, and manuscript production. All authors approved the final version of the manuscript.

## Data Availability

Raw and processed sequencing files can be found on the Gene Expression Omnibus super series ().

## Code Availability

Scripts used for analysis in this manuscript can be found at (https://github.com/jamrute).

## Materials and Methods

### Ethical Approval for Human Specimens

This study complies with all ethical regulations associated with human tissue research. Acquisition of donor samples was approved by the Washington University Institutional Review Board (study no. 201104172). All samples were procured with informed consent from patients obtained by Washington University in St Louis School of Medicine. No compensation was provided for participation. Cardiac phenotyping was performed at the time of LVAD implantation and explanation. Data were re-analyzed from GSE226314 and all associated clinical variables can be found in the associated manuscript^6^ Supplementary Tables 1-3.

### Human Recovery-Gene Expression Correlation

To connect transcriptional changes with patient ejection fraction within the LVAD cohort we computed a Pearson correlation coefficient between expression of the gene and patient ejection fraction at the pseudobulk level: 1) Normalized counts were aggregated to the pseudobulk level within each cell type and 2) a Pearson correlation coefficient was computed between every gene and patient ejection fraction within each cell type. Finally, genes which had a p-value < 0.05 and Pearson correlation coefficient > 0.6 or Pearson correlation coefficient < −0.6 were deemed highly correlated. The number of highly correlated genes were tabulated and cell types of rank sorted. Genes which had a Pearson correlation coefficient > 0.6 were considered positively associated with function and genes which had a Pearson correlation coefficient < −0.6 were considered negatively associated with function.

Pseudobulk differential expression to isolate a recovery signature: Pseudobulk differential gene expression was performed using the DESeq2 package^98^. We used data and the methodology previously described in GSE226314^6^: Briefly, we used the QCed and annotated integrated object and subsetted for each cell type, raw counts extracted, raw counts were aggregated to the sample level, data normalized using a regularized log transform, and differential expression analysis between conditions of interested via DESeq2. For pseudobulk DE analysis, we made the following comparisons: 1) HF vs donor, 2) Recovery vs donor, and 3) Recovery vs HF. Genes were deemed statistically significant if adjusted p-value < 0.05 and absolute(log2FC) > 0.58. Statistically significant DE genes from comparisons 1) – 3) were then used to identify a cardiac recovery gene set.

### Human Gene Program Identification and Prioritization

To explore transcriptional programs underlying recovery in myeloid cell states, we first performed non-negative matrix factorization (NMF)^40,41^ to generate a biologically meaningful low-dimensional embedding. Briefly, SCTransform-normalized^47^ data from the macrophage Seurat object from GSE226314 was used to identify the top 3,000 variable genes, followed by NMF decomposition using 30 components (k = 30) via the runNMF function in the GeneNMF package. A maximum of 100 genes were used for programs. The resulting NMF embeddings were then used to compute UMAP projections (RunUMAP) and identify transcriptionally distinct clusters based on shared nearest neighbor graphs and Louvain community detection (FindNeighbors, FindClusters). To identify consensus transcriptional programs across samples, we implemented consensus NMF (cNMF) using the multiNMF function on individual sample-specific Seurat objects, testing ranks from k = 4 to 9. Meta-programs were derived using getMetaPrograms, which aggregates stable gene programs across runs based on coherence and explained variance. We retained meta-programs explaining at least 70% of the variance, filtered based on quality metrics (positive silhouette score), and compiled gene sets corresponding to the top-ranked meta-programs for downstream pathway analysis and interpretation.

Deep learning prioritization of gene programs: To quantify the ability of meta-program features to predict cardiac recovery, we applied the Augur^42^ framework for gene program prioritization. First, a custom assay was constructed in the Seurat object using gene scores from selected consensus meta-programs (1–25), which were previously derived from cNMF above. We then subset the Seurat object to include only pre-LVAD samples leveraging the paired follow-up clinical data of whether the patient recovered or remained a non-responder and performed Augur analysis using the calculate_auc function. Discrimination performance was quantified by the area under the curve (AUC) of a classifier trained within the macrophages, with feature importance scores extracted to prioritize gene programs.

### Single-cell enhancer-to-gene mapping and TF nomination

We used the scE2G*^Multiome^* model^43^ to predict enhancer–gene connections in macrophages from human heart failure 10x Multiome data (GSE218392^25^) (https://github.com/EngreitzLab/scE2G/tree/v1.2). Briefly, scE2G first defines a set of candidate element-gene pairs where candidate elements are derived from peaks called on the pseudobulk ATAC-seq data for all macrophages and are paired with genes with a transcription start site within 5 Mb. Next, it annotates each pair with 7 features calculated from the macrophage multiome data, including 1) a pseudobulk Activity-By-Contact (ABC) score, where 3D contact is estimated by an inverse function of genomic distance; 2) the Kendall correlation across single cells between element accessibility and gene expression; 3) whether the gene is “ubiquitously expressed,” and 4) several other measures of genomic distance and chromatin accessibility around the element and promoter. Then, the features are integrated to assign each candidate element-gene pair a score representing the likelihood that the element regulates expression of the gene. Regulatory enhancer–gene interactions were defined as element–gene pairs with a score greater than 0.177, which is as the score yielding 70% recall when evaluating predictions in K562 cells against CRISPRi-validated enhancer-gene pairs^99^. We then used the scE2G macrophage map and genes from the top 3 gene programs predictive of recovery to identify disease relevant regulatory elements to perform TF enrichment analysis. TF enrichment analysis was performed using DoRothEA or the ENCODE and ChEA Consensus TFs from ChIP-X in EnrichR on the list of marker genes for the recovery associated pathways and no background set was used.

### In silico TF perturbation

Gene regulatory network construction: CellOracle (v0.10.5)^100^ was used to perform GRN analysis in macrophages. Processed data was imported into scanpy as .h5ad files. Highly variable genes (3,000) were kept for GRN construction. A promoter DNA sequences base GRN was initialized for humans. Unscaled counts were used to generate the CellOracle object and KNN imputation was performed. Cell state specific GRN were then constructed and the processed GRN was saved for subsequent TF KO analysis. For *in silico RUNX1* KO simulation in macrophages, the processed oracle object and inferred GRNs were loaded. The GRN was then fit using ridge regression models for the simulation. To simulate a KO, the *RUNX1* expression was set to 0, Next, we calculated cell state transition probabilities which are visualized as vectors on a digitized grid. To establish a baseline developmental flow field, we use pseudotime analysis: Palantir^101^ was used to perform pseudotime analysis to generate the developmental vector field. Palantir was used to compute a PCA, diffusion map, and a multiscale low dimensional embedding computer with 5 eigenvectors. The Palantir simulation was executed with classical monocytes as the starting state, no terminal states specified, and 600-1600 waypoints used. Next, the Gradient_calculator function from the CellOracle library was used to calculate a developmental vector field on the digitized grid. As described in CellOracle, an inner product was calculated between the baseline vector field and the post-KO simulation vector field to assess perturbations in different cell states to understand which populations are enriched and depleted post *RUNX1* KO. All visualization parameters were used as per CellOracle recommendations.

### Animal Models

Animal studies were performed in compliance with guidelines set forth by the National Institutes of Health Office of Laboratory Animal Welfare and approved by the Washington University institutional animal care and use committee. Animals were housed in a controlled environment with a 12 h light–dark cycle, with free access to water and a standard chow diet. All mouse strains used in this study are on the C57BL/6 background. *Runx1*^fl/fl^ (JAX #008772) mice were crossed to the *Cx3cr1*^creERT2^ (JAX #020940) to generate *Runx1*^fl/fl^*Cx3cr1*^creERT2^ strain. *Brd4*^fl/fl^*Cx3cr1*^creERT2^ were also used at the Gladstone Institutes with *Brd4*^fl/fl^ used as controls^31^. Additionally, a previously generated line from the Alexanian Lab at the Gladstone Institutes was also used for anti-FLAG CUT&RUN experiments (*Brd4*^Flag/Flag^*Cx3cr1*^creERT2^)^31^. Experiments were performed on mice 8-16 weeks of age and individual experiments contained mice of similar ages in all experimental conditions. Similar numbers of male and female mice were used for experiments. For Cre recombination in *Cx3cr1*^creERT2^ the background mice were given i.p. injection of 60□mg□kg^−1^ of tamoxifen (Millipore Sigma, T5648) with a 5-day pulse and then every 3 days for the duration of the experiment (or as indicated). For baseline deletion of *Runx1*, 5 days of i.p. injection of 60□mg□kg^−1^ of tamoxifen (Millipore Sigma, T5648) was given. Controls for all experiments include Cre− littermate controls treated with tamoxifen.

### Flow cytometry for characterization of baseline deletion on myeloid lineage in the heart

Hearts were perfused with 10mL of cold PBS, weighed, minced, and digested for 40□min at 37□°C in 3mL of Dulbecco’s modified Eagle medium (DMEM) (Gibco, 11965-084) containing 167□µl of 4,500□U□mL^−1^ collagenase IV (Millipore Sigma, C5138), 75□µl of 2,400□U□mL^−1^ hyaluronidase I (Millipore Sigma, H3506), and 30□µl of 6,000□U□mL^−1^ DNAse I (Millipore Sigma, D4527). Enzymes were deactivated with Hanks’ balanced salt solution containing 2% FBS and 0.2% BSA and filtered through 70□µm strainers. Cells were incubated with ammonium-chloride-potassium (ACK) lysis buffer (Gibco, A10492-01) for 3□min at room temperature. Cells were washed with DMEM and resuspended in 100□µl of FACS buffer (PBS containing 2% FBS and 2□mM EDTA). Cells were incubated in a 1:200 dilution of fluorescence-conjugated monoclonal antibodies for 30□min at 4□°C. Samples were washed with 1mL of FACS buffer and resuspended in a final volume of 300□µl FACS buffer. Flow cytometry was performed using a Cytek Aurora with Cytek SpectroFlo software. For analysis of neutrophils, dendritic cells, monocytes, and macrophages in the heart, the antibodies used were BV421 anti-mouse CCR2 (Biolegend, 150605), BV605 anti-mouse Ly-6G (BioLegend, 127639), BV711 anti-mouse CD64 (BioLegend, 139311), BV785 anti-mouse/human CD11b (BioLegend, 101243), FITC anti-mouse Ly-6C (BioLegend, 128005), PerCP-Cyanine5.5 anti-mouse CD45 (BioLegend, 103132), PE anti-mouse XCR1 (BioLegend, 148203), APC anti-mouse CD11c (BioLegend, 117309), Alexa Fluor 700 anti-mouse CD172a (BioLegend, 144022), and APC-Cyanine7 anti-mouse I-A/I-E (BioLegend, 107627).

### Closed chest ischemia reperfusion injury

Mice were anesthetized, intubated and mechanically ventilated. The heart was exposed and a suture placed around the proximal left coronary artery. The suture was threaded through a 1□mm piece of polyethylene tubing to serve as the arterial occlude. Each end of the suture was exteriorized through the thorax. The skin was closed, and mice were given a 2-week recovery period before induction of ischemia. After 2□weeks, the animals were anesthetized and placed on an Indus Mouse Surgical Monitor system to accurately record electrocardiography during ischemia. Ischemia was induced after anesthetizing the animals. Tension was exerted on suture ends until ST-segment elevation was seen via electrocardiography. Following 90□min of ischemia time, tension was released, and the skin was then closed.

### Echocardiography

Mice were sedated with Avertin (0.005□ml□g^−1^), and two-dimensional and motion mode images were obtained in the long and short axis views 3, 8, and 12 weeks after IRI using the VisualSonics770 Echocardiography System. Left diastolic volume, left systolic volume, akinetic area and total LV area were measured using edge detection and tracking software (VivoLab). The EF was calculated as (left diastolic volume□−□left systolic volume)/left diastolic volume. The akinetic percentage was calculated as the akinetic area/total LV area.

### Histology

Eight□weeks after IRI, mice were euthanized, and the hearts perfused with PBS. The hearts were fixed overnight with 4% paraformaldehdye (PFA) in PBS at 4□°C, cut into thirds on the transverse plane, placed into histology cassettes for paraffin embedding and dehydrated in 70% ethanol in water. The hearts were paraffin embedded and cut into 4□µm sections and mounted on positively charged slides. Tissues were stained by Gomori’s trichrome staining (Richard-Allan Scientific, 87020). Trichrome staining was imaged using a Meyer PathScan Enabler IV. Measurement of infarct length and LV length was performed in Zen Blue, with data quantification normalizing the length of the infarct to the length of the LV.

### Nuclei Isolation Multiome/ICELL8 sample preparation

Twelve□weeks after IRI, mice were euthanized, and the hearts perfused with PBS. The hearts were removed and flash frozen for downstream nuclei isolation. Briefly, the 10x Genomics Nuclei Isolation Kit (10x Genomics, PN1000494) was used: 3 sham, 3 WT, and 3 *Runx1*-KO hearts were dissociated in separate Eppendorf tubes and then all dissociations were combined at the end into 3 separate samples (one for sham, WT, and *Runx1*-KO hearts). Half the nuclei were used for downstream Multiome preparation, protocol CG000338 from 10x Genomics was used for Chromium Next GEM Single Cell Multiome ATAC + Gene Expression. Briefly, following nuclei isolation, permeabilization was performed, followed by transposition, GEM generation and barcoding using ChipJ (10x Genomics; PN1000234), post-GEM clean up, pre-amplification PCR, cDNA amplification, library construction, and sequencing. Gene expression and ATAC libraires were sequenced to a read depth of 50,000 and 25,000 respectively on a NovaSeq X. Remaining nuclei were used for processing with the ICELL8 platform. Briefly: we followed the SMART-Seq Pro Application kit (Takara Biosciences) to prepare samples for dispense by the iCell8cx system and subsequent reverse transcription, cDNA amplification, and library generation. Libraries were sequenced on one full lane of NovaSeq X Plus.

### Multiome data processing

Raw fastq files were aligned to the mouse mm10 reference genome using CellRanger ARC (10x Genomics, v6.1). We performed integrated single-nucleus RNA and ATAC analysis using Seurat v4 and Signac. Fragment files and peak calls were obtained from 10x Genomics Cell Ranger ARC outputs from sham, WT, and *Runx1*-KO hearts. Genomic peak coordinates were converted to GRanges objects and merged to generate a unified set of non-overlapping peaks. Peaks exceeding 10 kb or shorter than 20 bp were excluded. Fragment files were then used to quantify accessibility across the unified peak set using FeatureMatrix. RNA and ATAC count matrices were stored as separate assays in Seurat objects with gene annotations derived from the EnsDb.Mmusculus.v79 database. We applied the following QC per nucleus, filtering based on the number of detected RNA (500 < nCount_RNA < 10,000) and ATAC counts (500 < nCount_ATAC < 10,000), nucleosome signal (nucleosome_signal < 1), TSS enrichment (TSS.enrichment > 1), and mitochondrial RNA content (percent.mt < 10). Cells passing QC were merged into a single multiome object, and peaks were re-called using MACS2. Peaks were filtered to retain standard chromosomes and exclude blacklisted regions before re-quantifying chromatin accessibility to create a new “peaks” assay. Each modality was independently normalized and dimensionally reduced. RNA data were normalized using SCTransform with regression of nCount_RNA and percent.mt, followed by PCA and UMAP (RunUMAP, 50 PCs). ATAC data were processed using TF-IDF normalization, SVD, and LSI-based UMAP embedding. To integrate modalities, we computed a joint neighbor graph using FindMultiModalNeighbors, which combines PCA and LSI embeddings, and constructed a joint UMAP for visualization. Clustering was performed on the weighted nearest-neighbor (WNN) graph (FindClusters, resolution = 0.2–0.5). Clusters were annotated based on canonical marker gene expression and chromatin accessibility profiles using FindAllMarkers.

### Polygenic recovery score

Using positive- and negative-recovery associated genes from GSE226314, we constructed positive and negative polygenic recovery scores (PRSs) per cell type at the pseudobulk level from the human recovery data using EF as a surrogate of systolic function. We then converted these gene sets into mouse orthologs and used the Seurat AddModuleFunction to construct gene set scores for positive- and negative-PRS per cell type to visualize as an aggregate heatmap in sham, WT, and *Runx1*-KO groups.

### Differential gene, peak, and motif enrichment analyses

To define cell type–specific regulatory programs between WT and Runx1-KO animals, we conducted differential RNA expression, chromatin accessibility, motif activity analysis, and peak-to-gene linkage using paired single-nucleus multiome data. For gene expression analysis, each major cell type (cardiomyocyte, fibroblast, myeloid, endothelium, endocardium) we subset and analyzed using Seurat’s FindAllMarkers function on SCTransform-normalized expression data. Differentially expressed genes (DEGs) were identified using a Wilcoxon rank-sum test, with results filtered by adjusted p-value and log2 fold change thresholds. Chromatin accessibility differences were computed for the same cell types using Signac’s FindMarkers on peak-level data. Logistic regression was used with read depth as a latent variable, and significant peaks were defined based on p-value (<0.005) and absolute log2 fold change (>0.25). Peaks with strong accessibility shifts (log2FC > 0.58 or < –0.58) were further annotated. To infer transcription factor (TF) activity, we applied chromVAR, using motif annotations from the JASPAR2020 vertebrate core collection. Motif deviations were calculated across all cells, and wilcoxauc (from presto) was used to identify motifs with differential activity between groups. To assess cis-regulatory interactions, we used Signac’s LinkPeaks function within each cell type to map differentially accessible peaks to nearby genes using a correlation-based approach. Only high-confidence peak-to-gene links were retained. The number of unique target genes linked to condition-specific peaks was used as a proxy for regulatory remodeling in each cell type. Genes linked to differentially accessible ATAC-seq peaks were then used for downstream pathway analysis with clusterProfiler R package. Genes linked to differentially accessible ATAC-seq peaks were converted to Entrez IDs and submitted to enrichGO, using the org.Mm.eg.db annotation database for Mus musculus. Enrichment was conducted against the Biological Process (BP) ontology, with the Benjamini-Hochberg method applied for multiple testing correction. A q-value cutoff of 0.05 was used to define significant GO terms.

### ICELL8 Data Analysis

iCellcx8 sequencing data was aligned to the mouse reference genome via Cogent AP (Takara Biosciences). Raw count matrices were loaded and processed separately for each condition, with gene identifiers standardized by extracting gene symbols from Ensembl-style IDs and resolving duplicated entries. Seurat objects were created with a minimum cell and gene threshold (min.cells = 5, min.features = 500), and merged into a unified object. Cells with extremely high transcript counts (>400,000 UMIs) were filtered out. The dataset was normalized and variance-stabilized using SCTransform, regressing out nCount_RNA and percent.mt. Dimensionality reduction was performed via PCA, followed by neighbor graph construction (FindNeighbors), clustering (FindClusters, resolutions 0.5–2.0), and UMAP embedding (RunUMAP). Clusters were annotated based on canonical marker expression into major cardiac cell types. Cell type–specific markers were identified using FindAllMarkers with default Wilcoxon rank-sum test parameters (min.pct = 0.1, logfc.threshold = 0.25), and used to validate cell type identity. To characterize key biological pathways in cardiomyocytes (CMs), we performed differential expression analysis using FindAllMarkers with default Wilcoxon rank-sum test parameters (min.pct = 0.1, logfc.threshold = 0.25) visualized curated gene sets in heatmaps grouped by sham, WT, and *Runx1*-KO. We then compared the sequencing resolution in cardiomyocytes from iCell8-derived versus 10x based snRNA-seq profiles. The two platforms were benchmarked by plotting violin plots for gene complexity, generating UpSet plots of detected gene overlap, and visualizing gene detection saturation curves using a bootstrapped accumulation function. These analyses demonstrated the enhanced gene detection sensitivity of iCell8 compared to snRNA-seq in cardiomyocytes.

### Single cell isolation for scRNA-seq from murine hearts

Eight□weeks after IRI, mice were euthanized, and the hearts perfused with PBS. Cells were isolated as before^22^. Briefly: tissues were minced using a razor blade on ice and transferred to a 15 mL conical tube containing 3mL DMEM with 170uL Collagenase IV (250U/mL final concentration), 35uL DNAse1 (60U/mL), and 75uL Hyaluronidase (60U/mL) and incubated at 37 °C for 45 min with agitation. After 45 min, the digestion reaction was quenched with 6 mL of HBB buffer (2% FBS and 0.2% BSA in HBSS), filtered through 40 µm filters into a 50 mL conical tube and transferred back into a 15 mL conical tube to obtain tighter pellets. Samples were then spun down at 4 °C, 1200rpm for 5 min and the supernatant was discarded. Pellet resuspended in 1 mL ACK Lysis buffer (Gibco A10492-01) and incubated at room temperature for 5 min followed by the addition of 9 mL DMEM and centrifugation (4 °C, 5 min, 1200 rpm). Supernatant was discarded and the pellet was resuspended in 2 mL FACS buffer (2% FBS and 2mM EDTA in calcium/magnesium free PBS) and centrifugation was repeated in above conditions and supernatant aspirated.

Samples were then incubated in PDPN (BioLegend, 127423, clone 8.1.1), CD140a (Biolegend, 135911, clone APA5), CD31 (Biolegend, 102409, clone 390), and CD45 (Biolegend, 103132, clone 30-F11) with a 1:200 dilution for 30 min. Solution was washed 3X with FACS buffer following same centrifugation as above and then resuspended in 500 uL of FACS buffer and 1 uL DAPI (BD Biosciences, 564907) and filtered into filter-top FACS tubes. Gating for fibroblasts was performed as follows: PDPN+, singlets, CD45-/CD31-, PDPN+/PDGFRa+. Gating for immune cells was performed as follows: singlets and CD45+. Cells were sorted into 300 uL cell resuspension buffer (0.04% BSA in PBS). Collected cells were centrifuged as above, pooled across mice, and resuspended in collection buffer to a target concentration of 1,000 cells/uL. Cells were counted on a hemocytometer before proceeding with the 10X Genomics protocol. cDNA construction and library preparation were performed for 3 libraries (sham, WT, *Runx1*-KO) using the 10xGenomics 3’v3.1 Gene expression libraires were sequenced to a read depth of 50,000 on a NovaSeq X. Flow cytometry data collection was performed on BD FACSChorus version 1.1 and software FACSDiva version 8. Flow cytometry data analysis was performed in FlowJo 10.8.2.

### Cosine Similarity Analysis

To assess transcriptional convergence between human cardiac recovery and *Runx1*-KO, we computed cell type–resolved cosine similarity between gene expression profiles in humans and mice. We used the human recovery data (GSE226314)^6^ and compared against murine single-nuclei RNA-seq profiles from WT and *Runx1*-KO hearts. Mouse raw count matrices were mapped from mouse to human orthologs using a custom gene conversion function. To enable cross-species comparison, both the human and mouse Seurat objects were subset to common cell types and genes (Cardiomyocyte, Fibroblast, Myeloid, Endothelium, and Endocardium). Each dataset was normalized using Seurat’s NormalizeData. Within each shared cell type, we extracted cells from the recovery condition in humans and the *Runx*-KO condition in mice. For each group, gene expression was averaged across all cells, and cosine similarity was calculated between human and mouse mean expression vectors using the proxy::simil function. The resulting similarity scores were ranked to determine which cell types exhibited the highest transcriptional alignment between species. We repeated this analysis to compare recovered humans to sham animals to get a baseline and regress out baseline effects. A mouse-human recovery shift score per cell type was then calculate as follows: 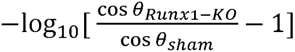 score indicates greater similarity between recovered *Runx1*-KO and human recovery.

### Single cell analysis of fibroblasts

Raw fastq files were aligned to the mouse mm10 reference genome using CellRanger and imported into Seurat objects. The datasets were merged and annotated with a condition metadata column. Quality control metrics, including number of detected genes (nFeature_RNA), total counts (nCount_RNA), and mitochondrial transcript percentage (percent.mt), were computed. Cells were retained if they had >200 and <7000 detected genes, <10% mitochondrial reads, and <30,000 UMIs. Normalization and variance stabilization were performed using SCTransform, with mitochondrial content and total RNA count regressed out. Principal component analysis (PCA) was computed using the top variable genes, and the top 30 PCs were used for dimensionality reduction via UMAP. Shared nearest neighbor (SNN) graphs were computed using FindNeighbors, and graph-based clustering was performed using the Louvain algorithm via FindClusters across a range of resolution parameters (0.1 to 0.5). Fibroblast cell states were annotated based on classical marker genes consistent with prior work^25,31,54^. To infer transcription factor (TF) activity, we applied chromVAR, using motif annotations from the JASPAR2020 vertebrate core collection. Motif deviations were calculated across all cells, and wilcoxauc (from presto) was used to identify motifs with differential activity between groups. To assess cis-regulatory interactions, we used Signac’s LinkPeaks function within each cell type to map differentially accessible peaks to nearby genes using a correlation-based approach. Only high-confidence peak-to-gene links were retained. The number of unique target genes linked to condition-specific peaks was used as a proxy for regulatory remodeling in each cell type. Genes linked to differentially accessible ATAC-seq peaks were then used for downstream pathway analysis with clusterProfiler R package. Genes linked to differentially accessible ATAC-seq peaks were converted to Entrez IDs and submitted to enrichGO, using the org.Mm.eg.db annotation database for Mus musculus. Enrichment was conducted against the Biological Process (BP) ontology, with the Benjamini-Hochberg method applied for multiple testing correction. A q-value cutoff of 0.05 was used to define significant GO terms.

### RNAscope In Situ Hybridization

Flash frozen LV samples were fixed for 24Lhr at 4□°C in 10% neutral buffered formalin, washed in 1× PBS, and embedded in paraffin. Paraffin-embedded sections were cut at an 8□μm thickness using a microtome. RNA In situ hybridization was performed using the RNAScope^102^ Multiplex Fluorescent Reagent kit v2 Assay, RNAScope 2.5 HD Detection Reagent as per the protocol – RED and RNAScope 2.5 HD Duplex Assay kits (Advanced Cell Diagnostics, ACDBio) using probes designed by Advanced Cell Diagnostics. Fluorescent images were collected using a Zeiss LSM 700 laser scanning confocal microscope. Chromogenic/brightfield images were acquired using a Zeiss Axioscan Z1 automated slide scanner. Image processing was performed using Zen Blue and Zen Black (Zeiss). Signal was quantified as number the of positive interstitial cells/total number of interstitial nuclei per 10x field.

### External In Vivo Dataset Analysis

We use publicly available data with annotations from the original manuscript^54^. Specifically, we extracted the myeloid cells. Using the human pseudobulk cardiac recovery genes, we calculated a gene signature for these genes in the mouse dataset and compared sham, TAC, TAC + JQ1, and TAC + JQ1 withdrawn.

### Mouse Model of TAC

*Brd4*^floxflox^*Cx3cr1*^CreERT2^ and *Brd4*^flag/flag^ mice underwent anesthesia with isoflurane, was mechanically ventilated (Harvard Apparatus) and then subjected to thoracotomy. During surgery, body temperature (37□°C) was maintained with the use of a temperature-controlled small-animal surgical table. Using 7-0 silk suture and a 25-gauge needle, the aortic arch was constricted between the left common carotid and the brachiocephalic as previously described^52^. For sham surgeries, thoracotomy was performed as above, and the aorta was surgically exposed without any further intervention. Mice were randomly assigned to either TAC or sham surgery groups.

### Langendorff Perfusion, Cell Isolation from post-TAC Cardiac Tissue

Cell isolation from *Brd4*^floxflox^*Cx3cr1*^CreERT2^ and *Brd4*^flag/flag^ TAC or Sham hearts were completed based of a previously described protocol with modifications. After the mouse was anesthetized, thoracotomy was performed and the heart was isolated and cannulated. Then using a Langendorff perfusion system (Radnoti, 120108EZ), the hearts were perfused with perfusion buffer (120.4□mM NaCl, 14.7□mM KCl, 0.6□mM KH_2_PO_4_, 0.6□mM Na_2_HPO_4_, 1.2□mM MgSO_4_, 10□mM Na-HEPES, 4.6□mM NaHCO_3_, 30□mM taurine, 10□mM 2,3-butanedione monoxime and 5.5□mM glucose, pHL7.0) for 5-10□min at 37□°C and then digested with digestion buffer (perfusion buffer with 300□U□ml^−1^ collagenase II (Worthington Biochemical LS004177) and 50□μM CaCl_2_) for another 10−min at 37□°C. After digestion, the atria and great vessels were removed, and the ventricular tissue was transferred to a dish and gently teased into small pieces in stop buffer (perfusion buffer with 10% fetal bovine serum) at 37□°C. After gently pipetting, the cell suspension was passed into a 50 mL conical tube through a 250□μM strainer and centrifuged at 30*g* for 3□min at 4°C. The supernatant containing most of the non-CMs was collected and centrifuged again at 30*g* for 3□min at at 4°C. The supernatant was again retained and filtered with a 70 μm cell strainer and centrifuged at 400*g* for 3□min at 4°C to eliminate debris. The non-CM cell pellet was then resuspended in 100□μl of FACS buffer (5% FBS, 0.01% NaN_3_ in 1× PBS) for downstream FACS application.

### Fluorescence Activated Cell Sorting from Isolated Cells of Cardiac Tissue post-TAC

The non-CM cell pellets obtained from the Langendorff perfusion and isolation described above from *Brd4*^flag/flag^ TAC and Sham hearts were resuspended in 100□μl of FACS buffer (5% FBS, 0.01% NaN_3_ in 1× PBS) and blocked with the Fc receptor antibody specific for FcγR III/II (BioLegend TruStain FcX PLUS anti-mouse CD16/32; 1:50 dilution) for 15□min. The block was quenched with 1 mL FACS buffer and cells were centrifuged at 500*g* for 5□min at at 4°C, resuspended in 100 μl of FACS buffer, and stained with a CX3CR1-APC antibody (BioLegend 149008, 1:100 dilution) for 30–40□min. After staining, cells were washed three times, each with 1 mL of FACS buffer and centrifuged at 500*g* for 5□min at at 4°C. After the final wash, the cells were resuspended in 400□μl of 5% FBS in 1X PBS, and transferred to a 5 mL polystyrene tube through a 35 μm filter cap. *Cx3cr1^+^* cells were sorted, gating for APC positive cells on the BD FACSAria II instrument. All flow cytometry analysis was performed using FlowJo (v.10.10). For the single-cell RNSA-seq performed in Sham and Day 7 post TAC in *Brd4*^floxflox^*Cx3cr1*^CreERT2^*and Brd4^f^*^loxflox^ animals, only immune cells were sorted, specifically cells were gated for mCD45-PB (Biolengend; 1:100), and against CD31-PE-Cy7 (Invitrogen; 1:50) and MEFSK4-APC (Miltenyi Biotec; 1:40). The staining and sorting strategies were the same as described above.

### Assessment of BRD4 chromatin occupancy by CUT&RUN in CX3CR1+ sorted cells

Whole hearts were obtained from *Brd4*^flag/flag^ animals that underwent sham or TAC surgeries and were subjected to Langendorff perfusion to obtain suspensions of cardiac cells. The non-CM population was purified by serial differential centrifugation and size-exclusion filtering steps followed by CX3CR1 antibody staining and sorting by flow cytometry as described above. Using the CUTANA ChIC/CUT&RUN Kit Version 3 (Epicypher) and an anti-Flag antibody (Sigma-Aldrich, M2, 1804, 0.5□μg per reaction, 1:100), CX3CR1+ cells were analyzed using CUT&RUN as previously described^103^ according to the manufacturer’s instructions with the included positive and negative controls. Purified fragmented DNA was quantified using the Qubit 2.0 fluorometer (Thermo Fisher Scientific) and the dsDNA HS Assay Kit. Paired-end Illumina sequencing libraries were prepared using the NEBNext Ultra II DNA Library Prep Kit (NEB E7645) and sequenced on the NextSeq 500 (Illumina) system. The nf-core/cutandrun (v.2.4.2; https://doi.org/10.5281/zenodo.5653535) pipelines were used to perform primary analysis within the Nextflow bioinformatics workflow manager (v.21.10.6) in conjunction with Singularity (v.2.6.0) using the following command: ‘nextflow run nf-core/cutandrun --genome GRCm38 -- input design_matrix.csv -profile singularity’. The pipelines perform adapter trimming, read alignment and filtering, normalized coverage track generation, peak calling and annotation relative to gene features, consensus peak set creation, and quality control and version reporting. All data were processed relative to the mouse (GCRm38, mm10) annotation.

### Generation of RAW 264.7 CRISPR Interference (CRISPRi) Line

RAW 264.7 mouse macrophage cell line was obtained from the American Type Culture Collection (ATCC) and cultured in high-glucose DMEM containing 10% FBS (SeraPrime F31016), 100□U□ml^−1^ penicillin-streptomycin (Gibco 15140122), 1X non-essential amino acid solution (Gibco 11140050) and 1□mM sodium pyruvate (Gibco 11360070) at 37□°C in a humidified incubator with 5% CO_2_. We generated a RAW 264.7 CRISPRi control line using a modified pHR-SFFV-KRAB-dCas9-mCherry vector (Addgene: 60954) that replaced the SFFV promoter with the EF1α promoter (pHR-EF1α-KRAB-dCas9-mCherry). To generate lentiviral particles, 3×10^6^ HEK-293T cells were seeded on a 100mm dish one day prior to transfection and cultured in 10 mL of supplemented high-glucose DMEM media. On the day of transfection, the old media was replaced with 10 mL of fresh media. This is followed by co-transfection of 5 μg of the desired lentiviral vector with 2.5 μg of the envelope protein vector pMD2.G (Addgene:12258) and 2.5 μg of the packaging vector psPAX2 (Addgene: 12260) using 59 μl of FUGENE HD transfection reagent (Promega E2312, San Luis Obispo, CA, USA) following manufacturer’s instruction. Supernatant was collected 48 hours post-transfection and filtered through 45 μm PVDF syringe filters (Thermo Fisher Scientific) to remove cell debris contamination. RAW 264.7 cells, which were seeded in one well of a 6-well plate, were then transduced with 2 mL of the lentiviral KRAB-dCas9-mCherry supernatant for 24 hours. We then harvested the RAW 264.7 cells 72 hours post-transduction and single-cell sorted mCherry positive cells into 96-well plates using flow cytometry (BD Aria II) to generate clonal lines expressing CRISPRi machinery. dCas9 expression was confirmed with qPCR and western blot and the RAW 264.7 line with the highest dCas9 expression was used in subsequent experiments. We further verified dCas9-Krab functionality by transducing the CRISPRi line with gRNA vectors targeting the transcription start site (TSS) of *Il1b*^104^ (see ‘CRISPRi guide RNA’ table) and assessed knockdown efficiency with qPCR.

### CRISPR interference (CRISPRi) for *Runx1* Enhancer Repression

CRISPRi was used for the targeted repression of *Runx1* putative enhancers in a RAW 264.7 macrophage CRISPRi line. The gRNAs targeting putative enhancers of *Runx1* were designed by the program ChopChop (http://chopchop.cbu.uib.no). We chose 3 gRNAs (see ‘CRISPRi guide RNA’ table) for every putative enhancer element targeting center of the peak and each side of the peak. We prioritized the ChopChop ranking for selecting the best guides with high efficiency (>50%), low self-complementary and low number of mis-match events. For constructing the gRNA lentiviral vector, we replaced the puromycin resistance gene with a hygromycin resistance gene in the pU6-sgRNA-EF1α-puro-T2A-BFP vector (Addgene: 60955), generating pU6-sgRNA-EF1α-Hyg-T2A-BFP. Pairs of gRNA oligos (see ‘CRISPRi guide RNAs table’) with 5’ (Sense oligo: 5’-TTG; and Antisense oligo: 5’-TTAGCTCTTAAAC) and 3’ (Sense oligo: 3’-GTTTAAGAGC; and Antisense oligo: 3’-CAACAAG) overhangs were synthesized (IDT, Coralville, IA, USA), annealed, and phosphorylated by T4 Kinase (New England Biolabs) and sub-cloned into BstXI and BlpI double-digested pU6-sgRNA-EF1α-Hyg-T2A-BFP by T4 ligase mediated ligation. The constructs were sequence verified (Quintara Bio, Berkeley, CA, USA). The RAW 264.7 CRISPRi clone was transduced with a mix of 0.5 mL of each of the three gRNAs viral supernatants (one gRNA targeting the center of the enhancer peak, the other two targeting the sides of the peak) for targeted repression of the putative enhancer. The gRNA viral supernatant was generated as described above. Pure polyclonal populations of cells harboring targeting gRNAs were selected with hygromycin (Thermo Fisher Scientific 10687010) treatment at 200 μg/ml for 10 days and repression efficiency was assessed by qPCR.

### Myeloid cell state analyses

Raw fastq files were aligned to the mouse mm10 reference genome using CellRanger and imported into Seurat objects. The datasets were merged and annotated with a condition metadata column. Cells were filtered to retain those with >200 and <8000 detected genes, <10% mitochondrial reads, and <30,000 UMIs. Mitochondrial percentage and UMI count were regressed out during normalization with SCTransform. Dimensionality reduction was performed using PCA, followed by UMAP embedding based on the top 50 PCs. Graph-based clustering was applied using the Louvain algorithm across multiple resolutions with harmony integration. Subsequent UMAP and clustering refined immune cell annotations into major immune cell types. A high-resolution subset of monocytes and macrophages was extracted and reclustered to reveal finer cell states. Marker genes were identified using FindAllMarkers, and clusters were annotated manually based on expression of canonical markers. Differential gene expression (DGE) was visualized using DotPlot and AverageExpression heatmaps. To infer transcription factor (TF) activity, we applied chromVAR, using motif annotations from the JASPAR2020 vertebrate core collection. Motif deviations were calculated across all cells, and wilcoxauc (from presto) was used to identify motifs with differential activity between groups. Transcription factor activity was inferred by integrating DoRothEA-derived mouse regulons. Target genes of downregulated motifs in *Runx1*-KO macrophages were compiled into a regulon module and scored using AddModuleScore.

### Assessing RUNX1 Chromatin Occupancy in RAW 264.7 Cells

CUT&RUN analysis was completed with some modifications as previously described^105^ for transcription factor enrichment on harvested RAW 264.7 cells using the CUTANA ChIC/CUT&RUN Kit Version 5 (Epicypher). In brief, 1×10^6^ cells per immunoprecipitation were resuspended in 1X PBS with 1mM EDTA and 0.5mM EGTA and fixed with 0.1% formaldehyde for 1 min then quenched with 125 mM glycine for 1 min at room temperature. Cells were then bound to 15 μL of activated ConA beads and permeabilized according to the kit manufacturer’s protocol. Immunoprecipitation was performed with 0.5 μg of anti-RUNX1 antibody (Abcam ab220117) and the provided positive and negative control and samples were incubated overnight on a nutator at 4°C. Samples were then washed and incubated on a nutator for 1 hour at 4°C after adding pAG-MNase. Chromatin cleavage was done according to manufacturer’s protocol and the reaction was stopped using a 2X Stop buffer comprised of 400mM NaCl, 20mM EDTA, 4mM EGTA, 0.02% Digitonin, 100ug/mL RNase, 50ug/mL Glycoblue, and 0.5ng/μL Epicypher Spike-in *E.coli* DNA. The chromatin was then released with a 37°C incubation for 30 minutes and the DNA was purified via phenol-chloroform-isoamyl alcohol extraction and ethanol precipitation. Sequencing libraries were generated with the NEBNext Ultra II DNA Library Kit (New England Biolabs (NEB) E7645) with transcription factor modifications^2^ and sequenced on the NextSeq 2000 (Illumina) system. The nf-core/cutandrun (v3.2.2-g6e1125d) were used to perform primary analysis within the Nextflow bioinformatics workflow manager (v.23.10.1 Build: 5891) in conjunction with Apptainer (v1.4.0-1.el8, formerly Singularity) using the following command: ‘nextflow run nf-core/cutandrun -profile singularity \ --gtf [gencode.vM35.annotation.gtf downloaded from Gencode] \ --fasta [GRCm39.primary_assembly.genome.fa downloaded from Gencode] \ --save_unaligned -- save_reference --input CnR_samplesheet.csv \ --outdir [destination path]’. The pipelines perform adapter trimming, read alignment and filtering, normalized coverage track generation, peak calling and annotation relative to gene features, consensus peak set creation, and quality control and version reporting. All data were processed relative to the mouse (GRCM39, vm35) annotation.

### Treatment of stimulated macrophages with Ro5-3335 in vitro

RAW264.7 cells (American Type Culture Collection (ATCC), TIB-71TM,) were grown in ATTC-formulated Dulbecco’s Modified Eagle Medium (30-2002, ATTC) supplemented with 10% fetal bovine serum, following ATCC guidelines. Cells were seeded in 12-well plates (Corning Inc, 3513) at a density of 3×10^5 cells/well and incubated for 24 hours before treatment. On day two, the spent media was gently aspirated from wells and replaced with fresh media, fresh media containing vehicle (DMSO) (100uM, Sigma-Aldrich, D1435-500ML), LPS (100ng/mL; Sigma, L4391), and/or Ro5-3335 (1uM or 100uM; MedChemExpress, HY-108470). After an additional 24 hours, supernatant from each well was collected and mouse interleukin-1b (IL-1b) and interleukin-6 (IL-6) protein levels were measured using MSD analysis (Meso Scale Discovery, MSD-V-PLEX Proinflammatory Panel 1 #K15048D-2). Cells were then washed with cold Dulbecco’s Phosphate Buffered Saline (GIBCO, 14190-144), RNA isolated using a RNeasy Mini Kit (Qiagen, 250102), and reverse transcription conducted using The High-Capacity cDNA Reverse Transcription Kits (ThermoFisher, 4368814). Digital PCR (dPCR) was performed using a QIAcuity Eight Platform Digital PCR System (Qiagen, 911056). TaqManTM gene expression assays: Mm00434228_m1 and Mm00446190_m1 (Thermo Fisher ScientificTM) for mouse Il-1b and mouse Il-6, respectively, and PrimeTimeTM qPCR probe, IDT Mm.PT.39a.22214839 (Integrated DNA Technologies/IDT) for mouse TATA-box binding protein (Tbp).

### In vivo Ro5-3335 administration

Wild type C57BL/6 animals at 8-weeks of age underwent IRI and subsequent echocardiography at 3-weeks post-IRI as described above. Ro5-3335 was formulated as follows: 100 mgs of Ro5-3335 were resuspended in 0.5 mL NMP (Sigma Aldrich, 328634-1L) and 2.5 mL PEG400 (Merck, 8.17003.500) and prepared fresh each week termed solution A. Solution A was then mixed with 1.75 mL of HPMC in citrate buffer pH 4. Animals were treated every other day at 40 mg/kg subcutaneously. Echocardiography was performed at 12-weeks post-IRI as described above in untreated and Ro5-3335 treated animals.

### Statistics and Reproducibility

No sample size calculations were performed. Sample size was governed by tissue availability and input tissue mass was based on ability to recover sufficient cells or nuclei. No samples were excluded. In vivo experiments were replicated at-least once for a total of two batches and data was pooled for downstream analysis (all attempts at replication were successful). For in vivo studies male and female mice were used and randomized to sham and injury groups and littermate controls used in genetic studies. Individuals performing surgeries and echocardiography measurements were blinded to the sample groups.

### CRISPRi guide RNA

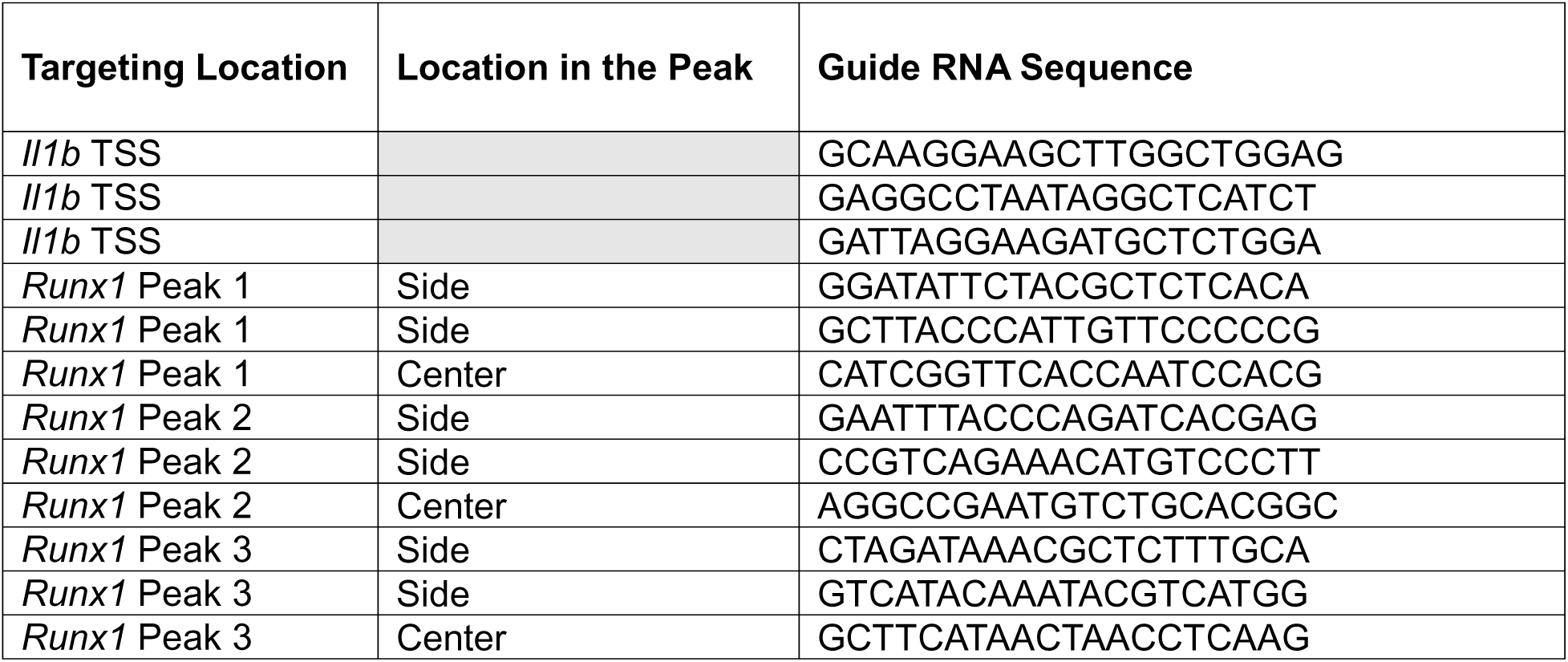

---

**Supplementary Figure 1.**
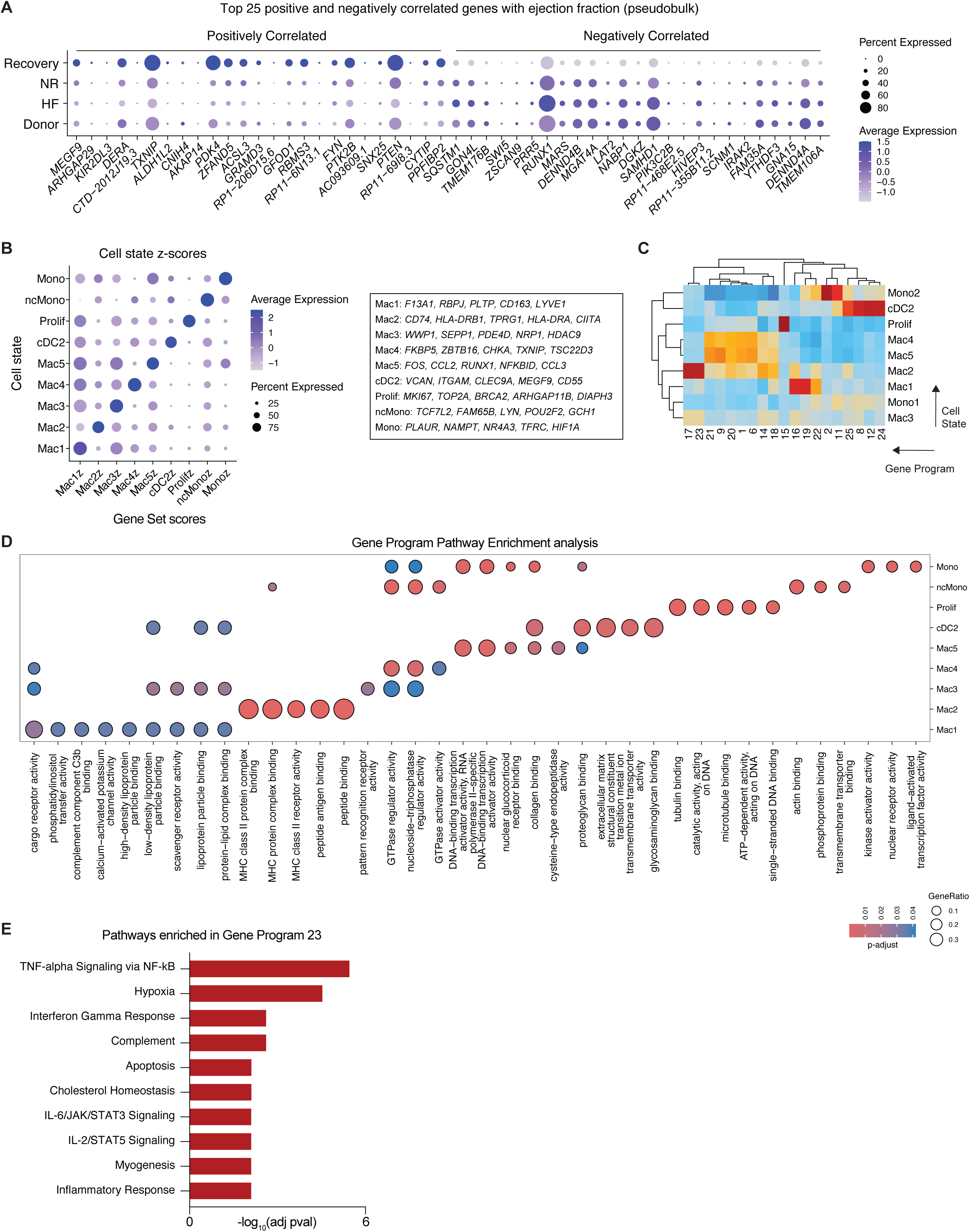
Macrophage gene programs in human recovery. (A) Dotplot of top 25 positively correlated (increase expression with ejection fraction) and 25 negatively correlated (decrease expression with ejection fraction) genes in macrophages in donor, failing, non-responders after mechanical unloading, and recovered human hearts. (B) Dot plot of gene set z-scores for top marker genes in each monocyte/macrophage cell state (left) with genes defined (right). (C) Heatmap of gene program z-scores in each monocyte and macrophage cluster. (D) Pathway enrichment analysis for marker genes in monocyte and macrophage cell states. The gene ratio is the proportion of genes from the pathway set which are within the cell state marker list. (E) Pathway enrichment analysis for gene program 23. The adjusted p-value in (D) and (E) is computed using the Benjamini-Hochberg method for correction for multiple hypothesis testing.

**Supplementary Figure 2.**
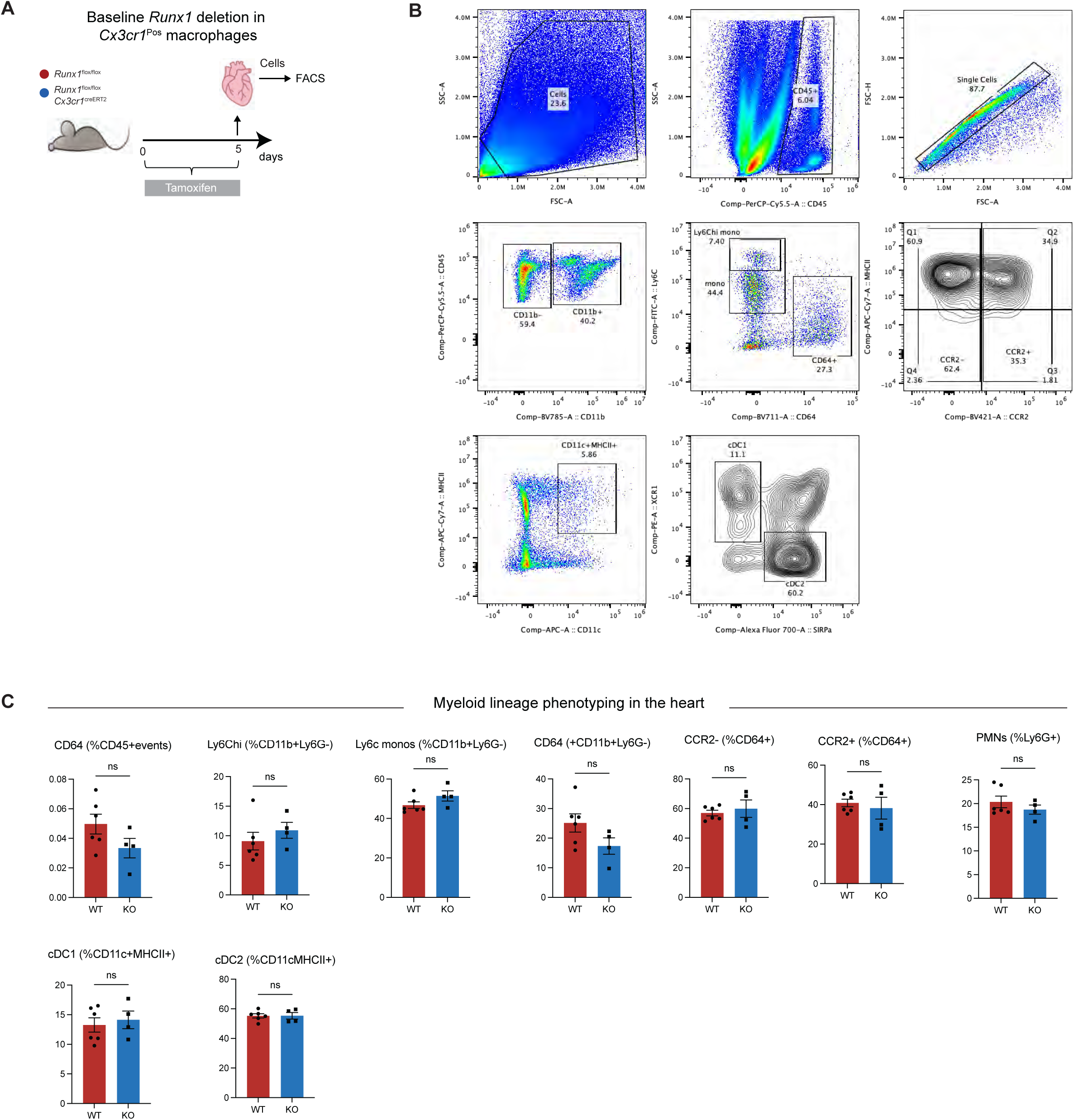
Baseline deletion of *Runx1* in cardiac macrophages does not impact the myeloid landscape in the heart. (A) Study design for targeted deletion of *Runx1* in cardiac macrophages at baseline with single cell isolation for flow cytometry in IRI and *Runx1*-KO animals. (B) Representative gating scheme to define the myeloid lineage subsets from the mouse heart. (C) Quantification of myeloid lineage cell states in IRI and *Runx1*-KO animals in the un-injured heart.

**Supplementary Figure 3.**
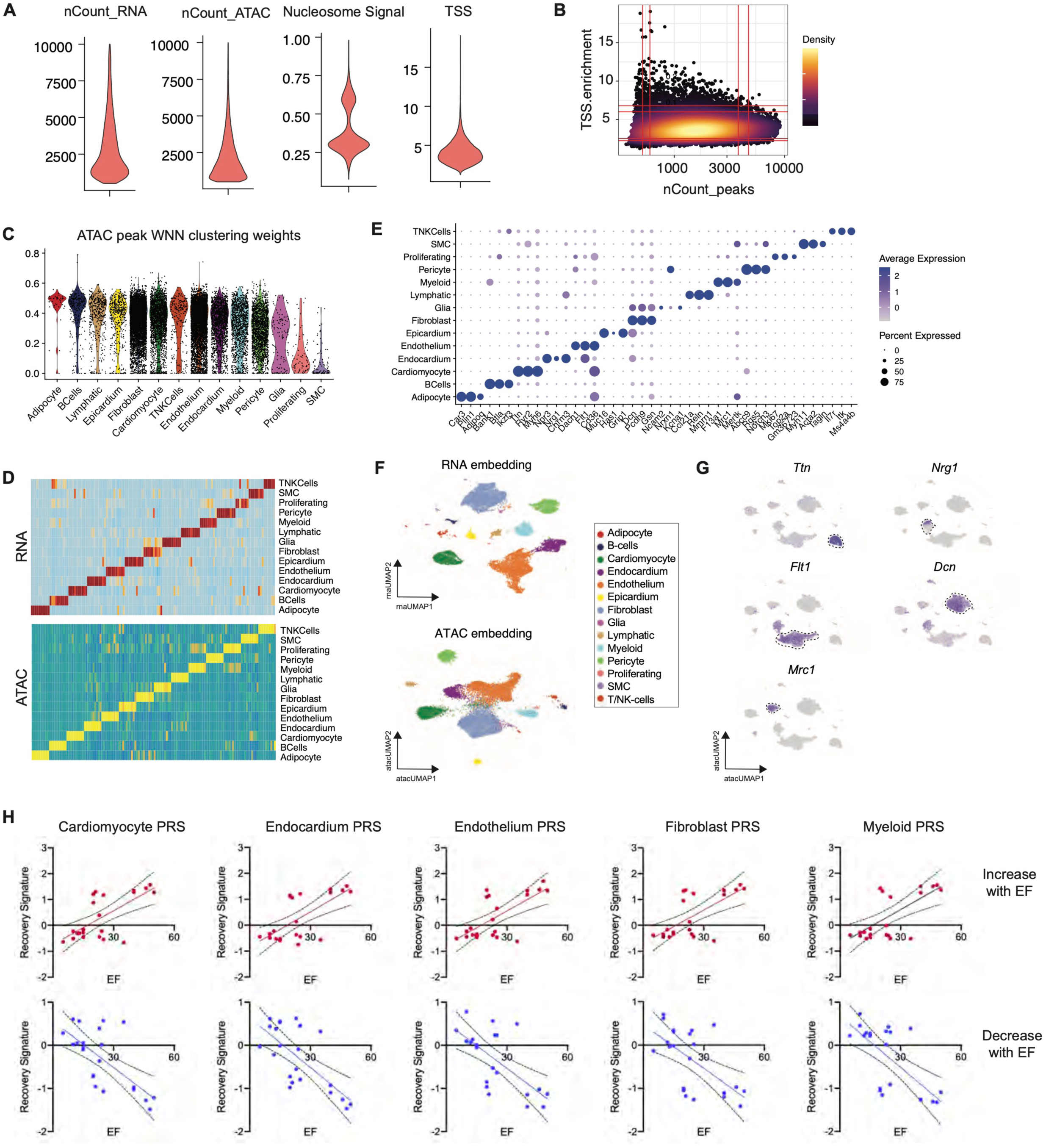
Quality control and clustering for Multiome data from sham, WT, and *Runx1*-KO. (A) Quality control metrics for number of RNA counts, number of genes, and mitochondrial RNA percent per nuclei. (B) Density plot for TSS enrichment and number of counts for ATAC peaks. (C) WNN clustering weights for ATAC data from Multiome data. Data was clustering using weighted gene expression and chromatin information. A higher value indicates more weight for chromatin used in clustering relative to gene expression. (D) Heatmap of differentially expressed genes and chromatin peaks by cell type. (E) Dot plot of canonical marker genes for annotated cell types from Multiome data. (F) UMAP embedding using only gene expression (top) or chromatin accessibility (bottom) data colored by cell type annotation. (G) Expression for *Tnt* (cardiomyocyte), *Nrg1* (endocardium), *Flt1* (endothelium), *Dcn* (fibroblast), and *Mrc1* (macrophage) to support cell type annotations. (H)

**Supplementary Figure 4.**
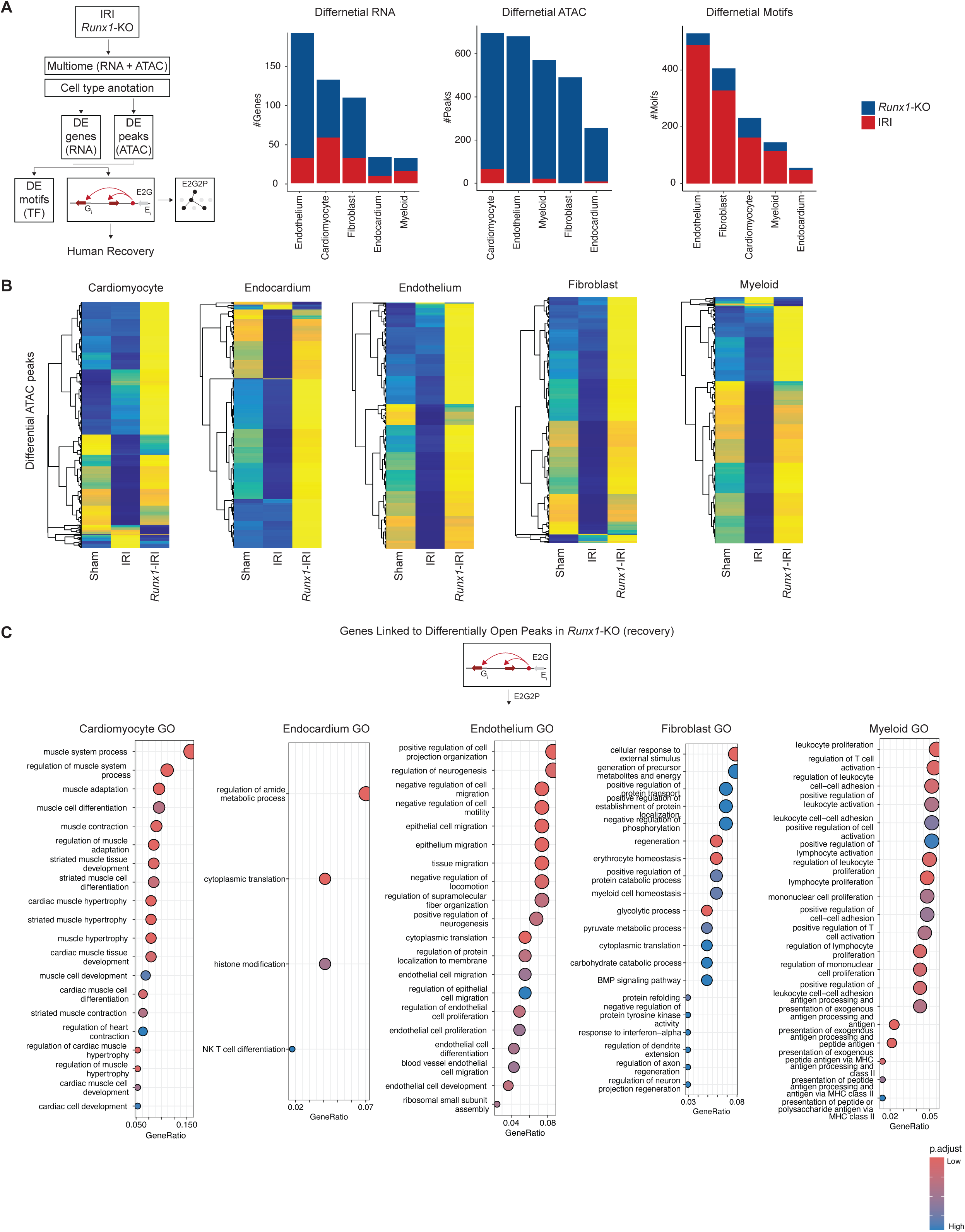
Cell specific differential gene expression, chromatin, and motif analysis in WT and *Runx1*-KO Multiome (RNA + ATAC). (A) Schematic of computational framework for integrative analysis of Multiome (RNA + ATAC) data (left) and tabulated number of differential genes, peaks, and associated motifs by cell type between WT and *Runx1*-KO animals 12-weeks post-IRI (right). (B) Heatmap of differential ATAC-peaks by cell type and grouped by sham, WT, and *Runx1*-KO. (C) Pathway enrichment analysis for genes linked to differentially open peaks in *Runx1*-KO relative to WT. E2G2P provides a framework to map regulatory changes to downstream transcriptional changes and associated functional pathways. The adjusted p-value is computed using the Benjamini-Hochberg method for correction for multiple hypothesis testing. The gene ratio is the proportion of genes from the pathway set which are within the cell state marker list.

**Supplementary Figure 5.**
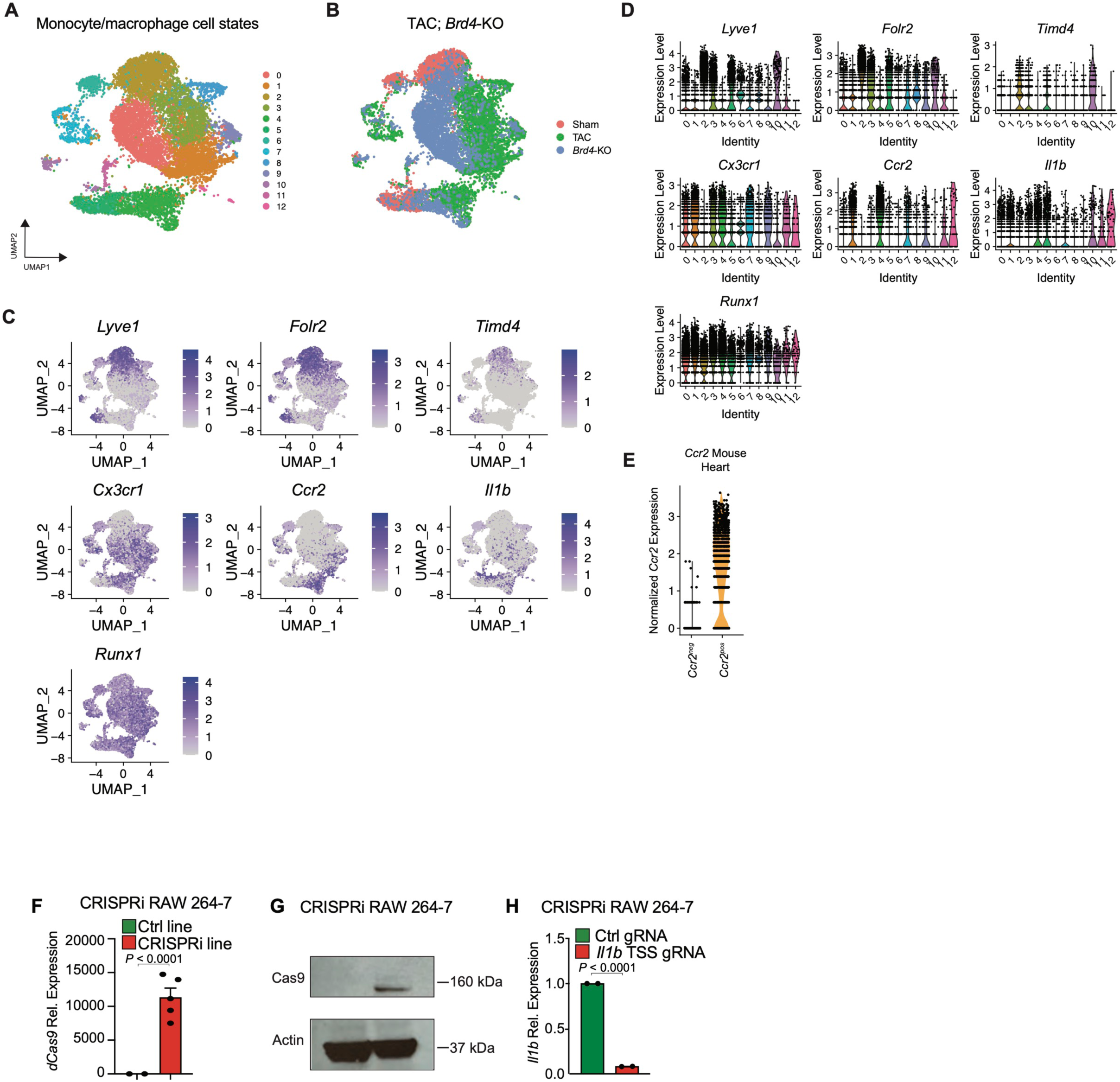
BRD4 regulation of *RUNX1* in cardiac macrophages. UMAP embedding of monocyte and macrophage cell states in *Brd4*-KO and WT animals post-TAC colored by cell state (A) and genotype/injury status (B). Expression of canonical macrophage genes on UMAP embedding (C) and represented as violin lots by cell state (D). (E) *Ccr2* expression violin plot split by *Ccr2*^neg^ and *Ccr2*^pos^ cell states. (F) Relative *dCas9* expression in control and CRISPRi RAW 264.7 macrophages. A higher relative expression in the CRISPRi line confirms *dCas9* insertion. (G) Western blot of protein levels for Cas9 in control and CRISPRi RAW 264.7 macrophages with Actin (house-keeping protein) as a reference. (H) qPCR for *Il1b* gene expression with a control non-targeting control and CRISPRi RAW 264.7 macrophages transfected with an *Il1b* TSS gRNA. A knockdown confirms active repression of targeted gene validating the CRISPRi line.

**Supplementary Figure 6.**
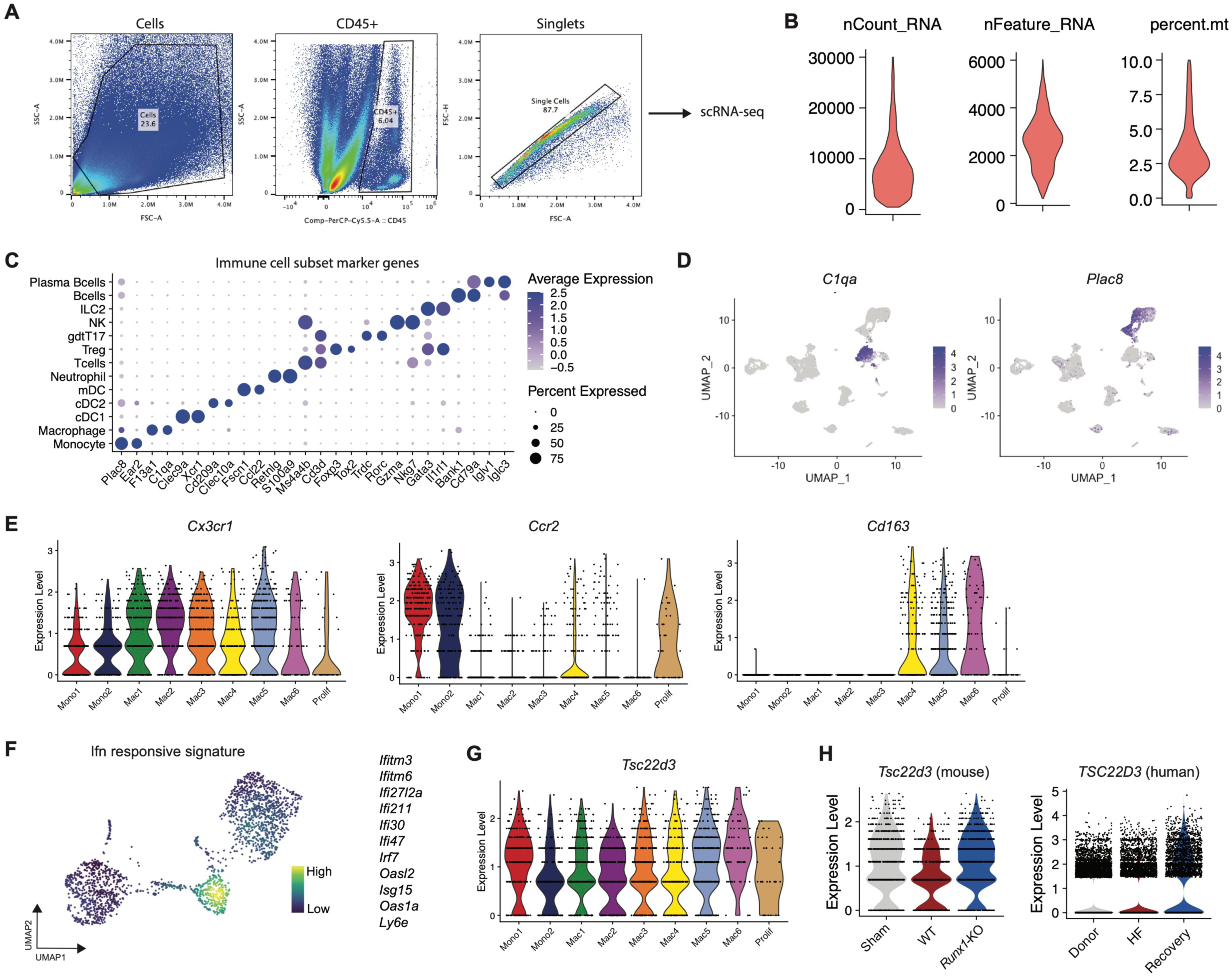
Monocyte and macrophage cell states in WT and *Runx1*-KO 8-weeks post-IRI. (A) Flow cytometry gating scheme for isolation of immune cells for single cell RNA-sequencing. (B) Quality control metrics for number of RNA counts, number of genes, and mitochondrial RNA percent per cell. (C) Dot plot of canonical marker genes for annotated immune cell types. (D) Gene expression for *C1qa* and *Plac8* to identify monocyte and macrophage cell types within UMAP embedding. (E) Violin plots for *Cx3cr1*, *Ccr2*, and *Cd163* expression grouped by monocyte/macrophage cell state. *Cx3cr1* marks all cardiac macrophages; *Ccr2* marks monocyte derived cells; *Cd163* marks tissue resident macrophages. (F) Gene set score density plot for type 1 interferon responsive genes (*Ifitm3, Ifitm6, Ifi27l2a, Ifi211, Ifi30, Ifi47, Irf7, Oasl2, Isg15, Oas1a,* and *Ly6e*). (G) Violin plot of *Tsc22d3* expression grouped by monocyte/macrophage cell state. (H) Violin plot for *Tsc22d3* expression in sham, WT, and *Runx1*-KO animals (left) and non-failing donors, heart failure samples, and recovered samples in human (right).

**Supplementary Figure 7.**
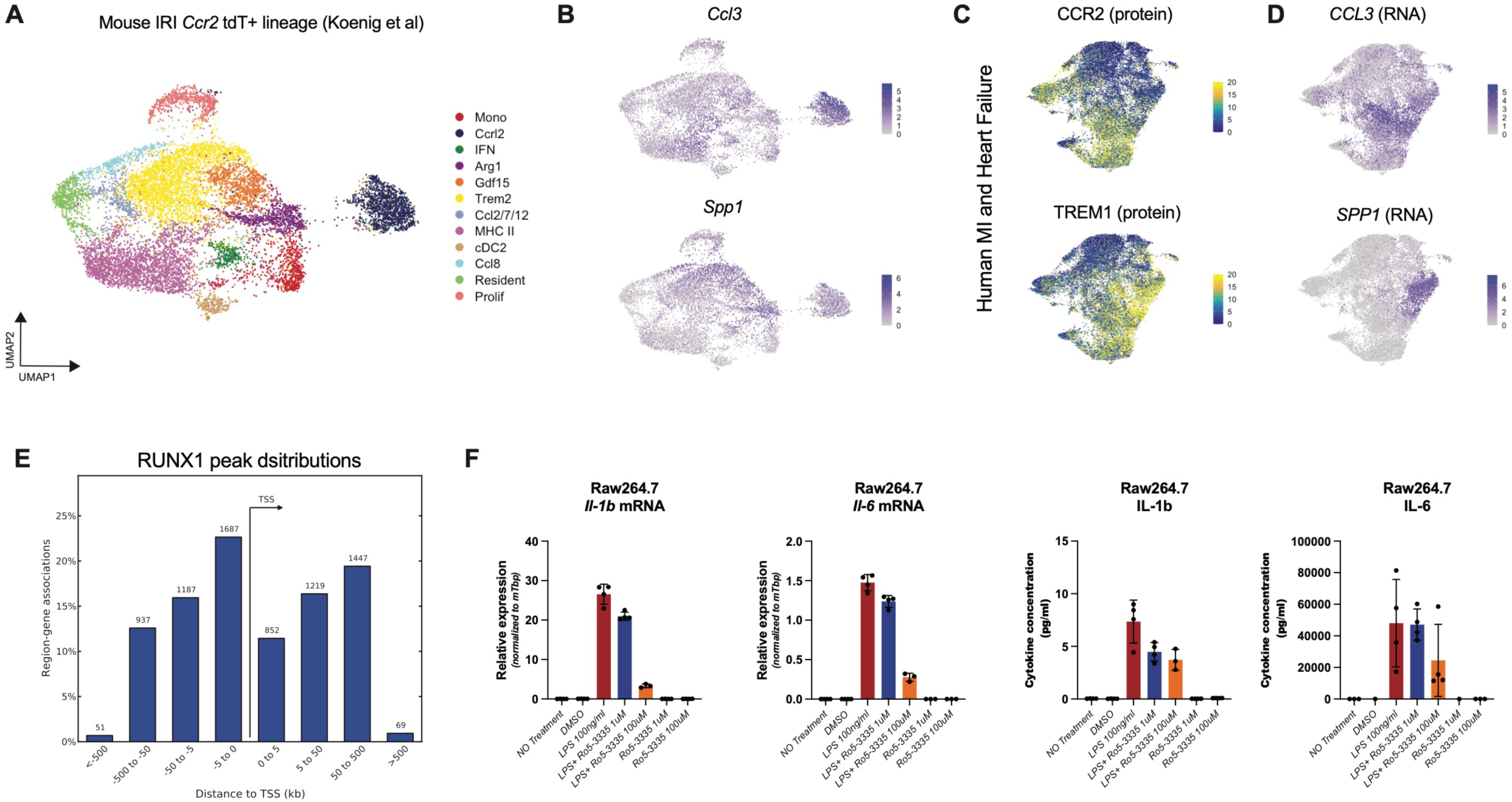
Convergence between mouse and human monocyte derived macrophages and RUNX1 regulatory network. (A) UMAP embedding plot for monocytes and macrophages post-IRI in *Ccr2*^CreERT2^ *Rosa26*-tdTomato animals. tdTomato positive cells were sorted for single cell sequencing therefore all cell states in the UMAP are monocyte derived populations. (B) *Ccl3* and *Spp1* expression in monocyte derived macrophages post-IRI. (C) CCR2 and TREM1 protein expression from human MI and HF CITE-seq in macrophages. (D) *CCL3* and *SPP1* expression from human MI and HF CITE-seq in macrophages. *CCL3* and *SPP1* co-localize with CCR2/TREM1 protein suggesting these chemokines are secreted by monocyte derived populations in human and mouse. (E) anti-RUNX1 CUT&RUN in RAW264.7 macrophages peak distributions showing maximal binding at the TSS with some distal gene regulation. (F) Dose dependent expression of *Il1b* and *Il6* RNA and protein levels in untreated, LPS treated, and LPS + Ro5-3335 treated RAW264.7 macrophages.

**Supplementary Figure 8.**
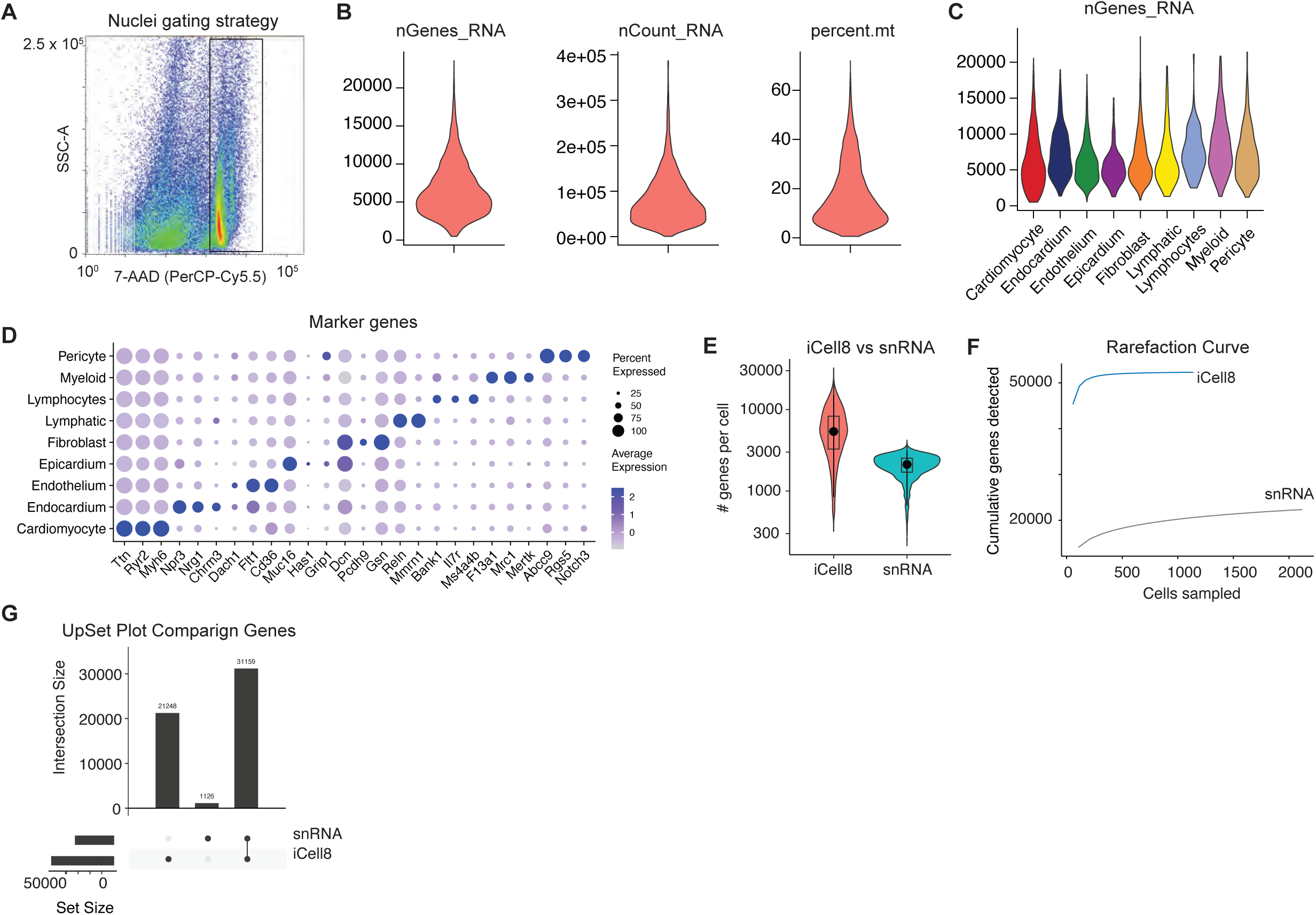
Ex vivo cardiomyocyte characterization using deep transcriptomics with ICELL8. (A) Nuclei gating strategy to sort 7-AAD+ nuclei from mouse hearts for ICELL8 profiling. (B) Quality control metrics for number of RNA counts, number of genes, and mitochondrial RNA percent per nuclei. (C) Number of genes detected per nuclei grouped by cell type. (D) Dot plot of canonical marker genes for annotated cell types from ICELL8 data. (E) Number of genes per nuclei in cardiomyocytes comparing ICELL8 versus single nuclei RNA from Multiome data. (F) Rarefaction curves comparing cumulative genes detected at different sampling rates in ICELL8 versus single nuclei RNA from Multiome data. Greater cumulative genes detected for the same number of nuclei sampled indicates greater data complexity. (G) UpSet plot comparing the number of unique genes detected in ICELL8 versus single nuclei RNA from Multiome data.

**Supplementary Figure 9.**
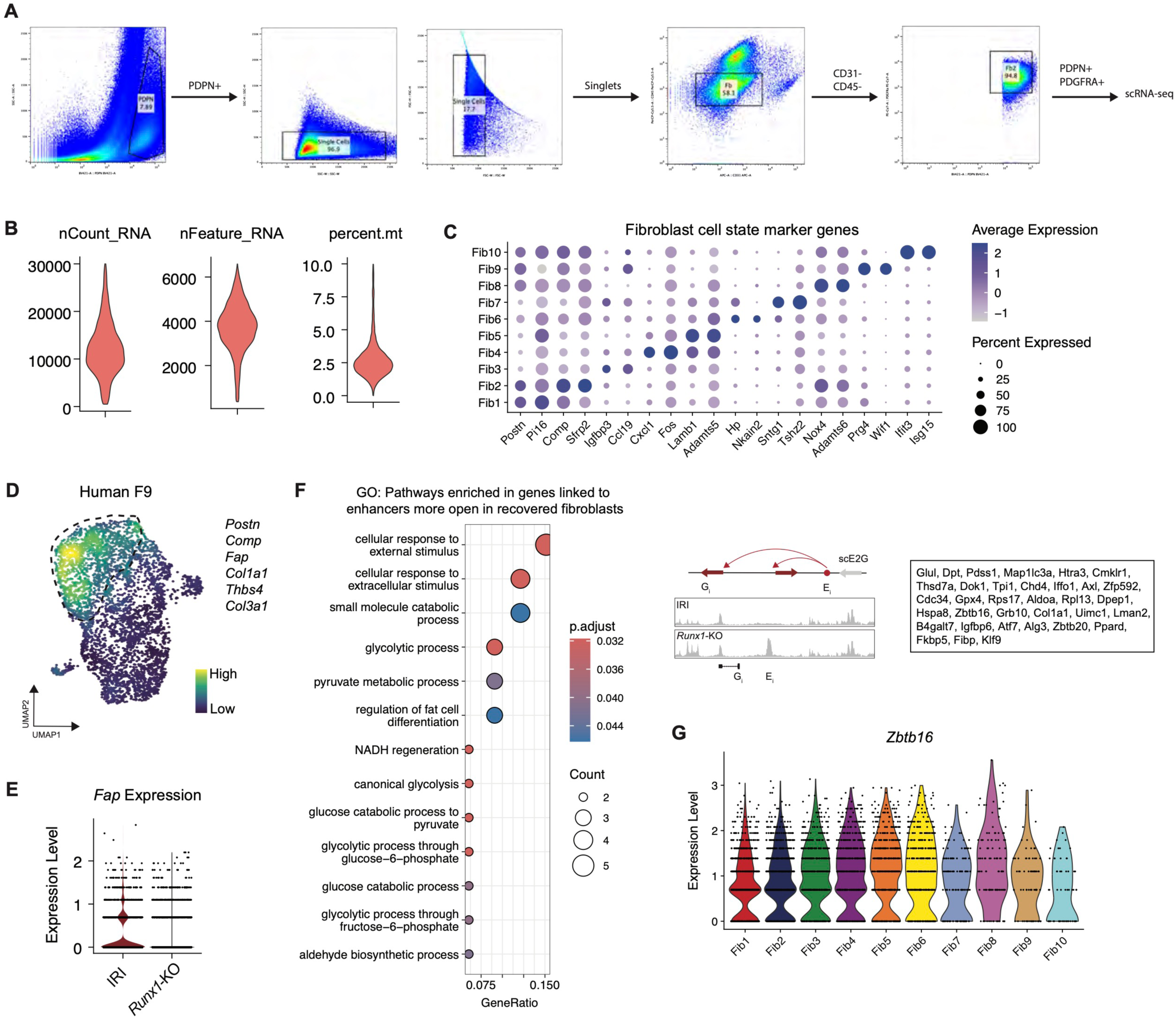
Quality control, cell state clustering, and pathway analysis in fibroblasts from WT and *Runx1*-KO animals 8-weeks post-IRI. (A) Flow cytometry gating scheme for isolation of fibroblasts for single cell RNA-sequencing. (B) Quality control metrics for number of RNA counts, number of genes, and mitochondrial RNA percent per cell. (C) Dot plot of canonical marker genes for annotated fibroblast cell states. (D) Human pro-fibrotic gene signature (*POSTN, COMP, FAP, COL1A1, THBS4,* and *COL3A1*)^25^ mapped on mouse cardiac fibroblasts. (E) Violin plot of *Fap* expression in WT and *Runx1*-KO fibroblast. (F) Gene ontology analysis for pathways enriched in genes linked to ATAC-seq peaks more open in *Runx1*-KO fibroblasts relative to WT fibroblasts (left) with schematic and genes (right). The adjusted p-value is computed using the Benjamini-Hochberg method for correction for multiple hypothesis testing. The gene ratio is the proportion of genes from the pathway set which are within the cell state marker list. (G) *Zbtb16* expression in cardiac fibroblasts grouped by cell state.

